# Structure-based design of small molecule inhibitors of the cagT4SS ATPase Cagα of Helicobacter pylori

**DOI:** 10.1101/2023.11.06.565890

**Authors:** Claire Morin, Vijay Tailor Verma, Tarun Arya, Bastien Casu, Eric Jolicoeur, Réjean Ruel, Anne Marinier, Jurgen Sygusch, Christian Baron

## Abstract

We here describe the structure-based design of small molecule inhibitors of the type IV secretion system of *Helicobacter pylori*. The secretion system is encoded by the□*cag*□pathogenicity island, and we chose Cagα, a hexameric ATPase and member of the family of VirB11-like proteins, as target for inhibitor design. We first solved the crystal structure of Cagα in a complex with the previously identified small molecule inhibitor 1G2. The molecule binds at the interface between two Cagα subunits and mutagenesis of the binding site identified Cagα residues F39 and R73 as critical for 1G2 binding. Based on the inhibitor binding site we synthesized 98 small molecule derivates of 1G2 to improve binding of the inhibitor. We used the production of interleukin-8 of gastric cancer cells during *H. pylori* infection to screen the potency of inhibitors and we identified five molecules (1G2_1313, 1G2_1338, 1G2_2886, 1G2_2889 and 1G2_2902) that have similar or higher potency than 1G2. Differential scanning fluorimetry suggested that these five molecules bind Cagα, and enzyme assays demonstrated that some are more potent ATPase inhibitors than 1G2. Finally, scanning electron microscopy revealed that 1G2 and its derivatives inhibit the assembly of T4SS-determined extracellular pili suggesting a mechanism for their anti-virulence effect.

## Introduction

*Helicobacter pylori* (*H. pylori*) is a Gram-negative bacterium isolated from gastric samples in the 1980s by Warren and Marshall. They demonstrated that the stomach is not sterile and that *H. pylori* causes inflammatory disease while colonizing the stomach (1). It was subsequently shown that chronic *H. pylori* colonization leads to gastric disease like ulcer, gastric adenocarcinoma and stomach cancer (2–4). *H. pylori* has been associated with humans for more than 1000 years and analysis of a strain from a human 5,300 years-old iceman showed a high degree of similarity with today’s *H. pylori* strains (5). Up to 50 to 90% of the human population is colonized by *H. pylori* and its prevalence depend on the geographic localization of populations in the world (6,7). *H. pylori* carry a lot of difference virulence factors for the colonization of the stomach, for the establishment of a niche between epithelial cells of the stomach and others that cause inflammatory responses in human cells (2,8,9). The main virulence factors are the urease that neutralises the acidic environment, the vacuolating cytotoxin A (VacA) and the type IV secretion system (T4SS) (8–10). The T4SS comprises 27 proteins coded by the cytotoxin-associated gene (*cag*) pathogenicity island (*cag*PAI). The T4SS in *H. pylori* is used for the injection of the oncoprotein CagA as well as for the transfer of peptidoglycan fragments (11–14). CagA interferes with different metabolic pathways in gastric epithelial cells, leading to depolarisation of the cell, an increase of cell mobility and the loss of tight junctions (8,15–17). The resulting disorganization of gastric cell structure contributes to gastric disease (18–20). Not all *H. pylori* carry a *cag*PAI, but it is a major factor contributing to gastric disease and cancer (20).

Antibiotic therapy is commonly used for the treatment of patients with gastric disease. The treatment comprises triple or quadruple therapy and often includes proton pump inhibitors and bismuth salts (21–25). Therapy with multiple antibiotics provokes a high selection pressure for antibiotic resistance. In addition, *H. pylori* is highly transformable, and it can acquire resistance that spreads in the environment. Over the last 10 years more than half of *H. pylori* patient isolates were found to carry at least one resistance gene against the first line antibiotics (26–28). In 2017 the World Health Organization published a report underlining the emergency of finding new antibiotics against *H. pylori* (29).

Considering the rise of antibiotic resistance, alternative therapeutic methods to fight bacterial infections have emerged that may not lead to strong selection pressure (30–33). Targeting virulence factors that are not essential for bacterial survival is one of the investigated approaches (34–37). *H. pylori* virulence factors such as the T4SS are not essential for its survival, but they are major contributors to severe gastric disease. The *cag*T4SS proteins are unique to *H. pylori* and inhibitors would therefore not target other bacteria of the microbiome. We have chosen the essential ATPase Cagα as a target (14,38,39). Cagα is a hexameric ATPase that localizes at the inner membrane and is a homolog of VirB11 proteins that are conserved and essential for all T4SS.

Previously, we have used fragment-based screening to identify small molecules that interact with Cagα and we identified 1G2 as a non-competitive inhibitor of its ATPase activity (37). Here, we characterized the binding site by X-ray crystallography and mutagenesis identified critical amino acid residues. Based on the information of the binding site we created a family of 1G2 derivates and some of them have interesting potential as inhibitors of T4SS activity and of extracellular assembly of Cag pili.

## Materials and Methods

### Cloning and mutagenesis

Cloning of Cagα was described by Arya *et al*, 2019 (37). The plasmid carrying the WT Cagα protein pHTCagα was amplified using the QuikChange II site-directed mutagenesis kit (Agilent) using 33 bp primers (suppl. Tab. 1) to introduce single codon mutations to change amino acid residues at the 1G2 binding site, followed by Sanger sequencing on an ABI 3730 sequencer.

### Expression and purification of Cagα

The expression and of purification Cagα wild type and mutant proteins was conducted as previously described by Arya *et al*, 2019 (37).

### Crystallisation and structure determination

Initial crystallization conditions were established using the MCSG screen from Anatrace (USA using 6 mg/ml of Cagα and 1 mM of 1G2 (1:10 ratio). Final crystals were grown at room temperature using the hanging drop vapour diffusion method in 100 mM Bis-Tris (pH 6.5) and 2mM ammonium sulfate. Drops containing 2 μL of protein-inhibitor-mixture (1:10 ratio) and 2 μL of reservoir solution were incubated for 2 weeks. Hexagonally-shaped crystals appeared after 7-10 days. The crystals were cryo-protected in 100 mM Tris*-*HCl buffer (pH 8.5), 2 M ammonium sulfate and 25% glycerol, flash frozen in liquid nitrogen and the data were collected at microfocus beamline F1 at the Cornell High Energy Synchrotron Source (CHESS). The intensity data was processed using the HKL2000(40) program (suppl. Tab. 2). The structure was solved by molecular replacement using the coordinates of PDB ID: 1G6O as search model. Refinement and modeling were performed using REFMAC and Coot (41,42). Final graphical figures and tables were generated using the Pymol-integrated Phenix software suite (43). The structure was published in the PDB (PDB code: 6BGE, https://www.rcsb.org/structure/6BGE).

### Docking of 1G2 and derivatives to Cagα

Autodoc Vina was used for binding energy calculations. The Cagα structure used for the binding from PDB 1G6O (RCSB PDB - 1G6O: CRYSTAL STRUCTURE OF THE HELICOBACTER PYLORI ATPASE, H. PYLORI0525, IN COMPLEX WITH ADP) was used for docking.

### Differential scanning fluorimetry (DSF)

To assess binding of 1G2 derivatives to Cagα and its mutant proteins, DSF was used in the presence of SYPRO orange as in Arya *et al*, 2019 (37).

### Synthesis of the 1G2 derivatives

The organic reagents are prepared as DMF solutions: Solution 1 : The heteroaryl halide building blocks solution was prepared by weighing 1.77 mmol in a 2 ml vial. DMF was added and the volume is adjusted to 400 µl. Solution 2 : The scaffold (methyl-4-hydoxybenzoate) was weighed (1259 mg; 8.27 mmol) in a 40 ml vial, the solid was dissolved with DMF and the volume adjusted to 29.4 ml. A 0.6-2 ml microwave vial was added with 161 mg (0.493 mmol; 2.5 eq.) of Cs_2_CO_3_, equipped with a stirbar, charged with the heteroaryl halide solution (100 µl; 2.25 eq.) followed by scaffold solution (700 µl; limiting reagent). The suspension was transferred to a Biotage Initiator microwave reactor to run the reactions at 180 °C for 1 h. The reaction was monitored by liquid chromatography mass spectrometry (LCMS). The reaction mixtures were filtered on a filtering plate (96 deep-well plate; PE Frit 25 mm; Long drip; 2 ml) using a MeOH/DMSO (2:1) mixture. Filtration was forced using HT4X Genevac for centrifugal filtration. The filtered mixtures were added to 50 ml of AcOH (pH adjusted <5) and purified on a reverse-phase Kinetex 5 µm C18 column 21.2 x 100 mm and was eluted with MeOH - Water - 0.1% AcOH. Gradient: Isocratic 25 or 30 % for 1.5 minutes then gradient to 100% MeOH over 8.5 minutes. Tubes containing the desired compounds were identified and placed on an HT6- Genevac evaporator. After removal of the solvent, the content of the tubes was transferred to pre-tared 4 ml vials using a (1:1) DCM:MeOH mixture. The vials were placed in a HT6 Genevac evaporator and the solvents removed under reduced pressure, followed by NMR and LCMS analysis.

### Enzyme activity assay

The ATPase activity was quantified using a malachite green binding assay (37). The 100□μL reaction mixtures contained 60 nM of enzyme, 25□mM HEPES (pH 7.5), 100□mM NaCl, 200□µM MgCl_2_ and 200 μM of 1G2 derivative. The reaction mixtures were incubated for 15□min at 30□°C. After incubation, ATP was added at 100 μM concentration and incubated again for 30 minutes at 30°C and then 40□μL of malachite green assay mixture was added. The formation of the blue phosphomolybdate-malachite green complex was in linear relation to the amount of released inorganic phosphate and measured at 610 nm. Color generated was estimated compared to series of standards.

### IC_50_Ldetermination

IC_50_□values were determined by incubating different concentrations of molecules (5–2,500□µM; from stocks of 100□mM) with enzyme in 25□mM HEPES (pH 7.5) and 100□mM NaCl and 200□µM of MgCl_2_. Mixtures were incubated with inhibitors for 15□min, followed by addition of ATP and incubation for 30□min at 37□°C. The reactions were stopped by addition of 40□µl malachite green solution and the inorganic phosphate released was determined at 610□nm. Data were plotted as 1/rate versus inhibitor concentration for each substrate concentration and a linear fit was calculated by non-linear regression using SigmaPlot (version 11.0).

### Bacterial strains, cell lines and culture conditions

*H. pylori* strain 26695 (ATCC700392) was used as a positive control, and the previously described Δ*cagV* (*H. pylori*0530) mutant recreated in our laboratory was used as negative control (37,44). All the strains were cultivated on Columbia agar base (BD) containing 10% (v/v) horse serum (Wisent Inc.), 5% (v/v) laked horse Blood (Wisent Inc.) with β-cyclodextrin (2 mg/mL), vancomycin (10 mg/ml) and amphotericin B (10 mg/ml). Chloramphenicol (34 mg/ml) was added in case of the Δ*cagV* strain to select for the chloramphenicol (*cam*) gene cassette used to disrupt the gene. For liquid cultures, brain heart infusion (BHI) media (Oxoid) was supplemented with 10% fetal bovine serum (FBS) and appropriate antibiotics. Bacteria were cultivated at 37°C under microaerophilic conditions (5% oxygen, 10% CO_2_). Gastric human cells (AGS CRL-1739) cells were grown at 37°C in F12K media (Wisent Inc.) with 10% (v/v) FBS (Wisent Inc.) in a 5% CO_2_-containing atmosphere.

### Cell proliferation Assay

AGS cells cultivated at 5x10^4^ cells/well density were incubated overnight with 100 μM of molecules at 37°C in a 5% CO_2_-containing atmosphere. The cell proliferation was evaluated using the Cell Proliferation Reagent WST-1 (CELLPRO-RO, Roche).

### Infection of AGS by H. pylori strains and IL-8 production

AGS cells cultivated at 7x10^5^ cells/well density in 6-well plates were infected for 4h or 0.3x10^5^ cell/well density in a microplate for 1h, with a culture of *H. pylori* at multiplicity of infection (MOI) 1:100. *H pylori* were first pre-incubated for 1h in F12K supplement with 10% FBS with or without 1G2 at concentration 200µM at 37°C in a 5% CO_2_-containing atmosphere. After 4h infection under microaerophilic conditions, supernatants were sampled and centrifuged at 15,000 *g* to remove cells and debris; the supernatants were conserved at -20°C. The level of IL-8 in cell culture supernatants was determined using a human IL-8 ELISA kit (Invitrogen).

### Sample preparation and western blotting

After 4h of infection, the cells were washed with phosphate-buffered saline (Wisent Inc.) and lysed in RIPA Buffer (50 mM Tris-HCl pH 8.0, 150 mM sodium chloride, 1.0% Igepal CA-630 NP-40, 0.5% sodium deoxycholate, 0.1% sodium dodecyl sulfate), complemented with a protease inhibitor cocktail for mammalian tissues (Sigma Aldrich) and a phosphatase inhibitor cocktail for tyrosine protein phosphatases, acid and alkaline phosphatases (Sigma Aldrich). The cells were harvested, incubated at 95°C for 5 min with SDS-PAGE sample buffer and centrifuged for 10 min at 10,000 rpm, followed by SDS-PAGE and western blotting. The production of proteins was assessed with monoclonal mouse anti-*Helicobacter pylori* CagA (HyTest Ltd.) and rabbit Cagα antiserum (Abcam). The actin was identified using the anti-β-Actin antibody (C4): m-IgG Fc BP-HRP (Santa Cruz, sc-528515). The secondary antibodies (rabbit and mouse) were purchased from Biorad and the HRP signal was developed using Clarity Western ECL Substrate (Biorad).

### Scanning electron microscopy

AGS cells were cultivated on round cover glasses (Fisherbrand) at 6x10^5^ cells/well density in 6-well plates. The cells were infected with *H. pylori* for 4h as explained above, washed with cold phosphate buffer (PB 0.1 M), fixed in 4% paraformaldehyde/0.1% glutaraldehyde for 30 min at 4°C, followed by another wash in PB. The samples were then incubated in osmium tetroxide 4% (0.1%) for 1 h at 4°C, followed by washing with PB. The samples were then dehydrated using a series of ethanol dilutions for 15 min each (30%, 50%, 70%, 80%, 90%, 95%, 100%, 100%), followed by drying using critical point dryer using the Leica EM CPD300. Then cells were then coated with 5 nm of carbon using the Leica EM ACE600. The sample were visualised using a Hitachi Regulus 8220 scanning electron Microscope. The length and number of T4SS pili were measured using ImageJ software (45).

### Statistical analysis

All the statistical analyses were performed using GraphPad Prism 9 (9.5.1). T-test and ANOVA (Kruskal-Wallis test and ordinary One-way) were used to analyse the number of pili and their size, respectively.

## Results

### X-ray analysis reveals the 1G2 binding site

In our previous work we identified 1G2 as a non-competitive inhibitor of Cagα and this molecule has interesting potential for development into an anti-virulence drug (37). As first step towards designing more effective 1G2 derivatives, we characterized the binding site by X-ray crystallography. Cagα was co-crystallized with 1G2, and we solved the X-ray structure of the Cagα-1G2 complex in the *P*6_3_22 crystal space group with two molecules in the asymmetric unit (Fig. 1A) to a resolution of 2.9 Å (suppl. Tab. 2). The structure was solved by molecular replacement using ADP bound Cagα (PDB code: 1G6O) as a search model. The overall structure of 1G2-bound Cagα (PDB code: 6BGE) is similar to that of the Cagα-ADP complex, but there are differences in interactions at the protein interface, and we identified the electron density of molecule 1G2 sandwiched between two Cagα molecules (Fig. 1A). The monomer structure of the Cagα-1G2 complex displays both N-terminal domain (NTD) and C-terminal domain (CTD) with nine α-helices labeled as α1 to α9 and 13 β-strands labeled as β1 to β3 (Fig. 1B). A structural overview from the top of the NTD reveals that 1G2 interacts with the NTD of both protein subunits (Fig. 1C). The 1G2 binding site is distinct from the active site to which ADP and the substrate analog ATP-γ-S bind (39,46). Molecule 1G2 binds to a hydrophobic pocket created by the interaction between the NTDs of two Cagα subunits and amino acids F68 and F39 make hydrophobic contacts with the two aromatic rings of the inhibitor. R73 and D69 are the amino acids involved in forming a polar contact with 1G2. R73 interacts with the pyridine ring via a hydrogen bond and the carboxylic group of 1G2 interacts with the backbone NH group of D69 forming a potential hydrogen bond (Fig. 1D). Sequence alignments with other Cagα /VirB11 homologs show that this site is not conserved suggesting that the molecule could likely be a specific inhibitor (suppl. Fig. 1).

**Figure 1:**
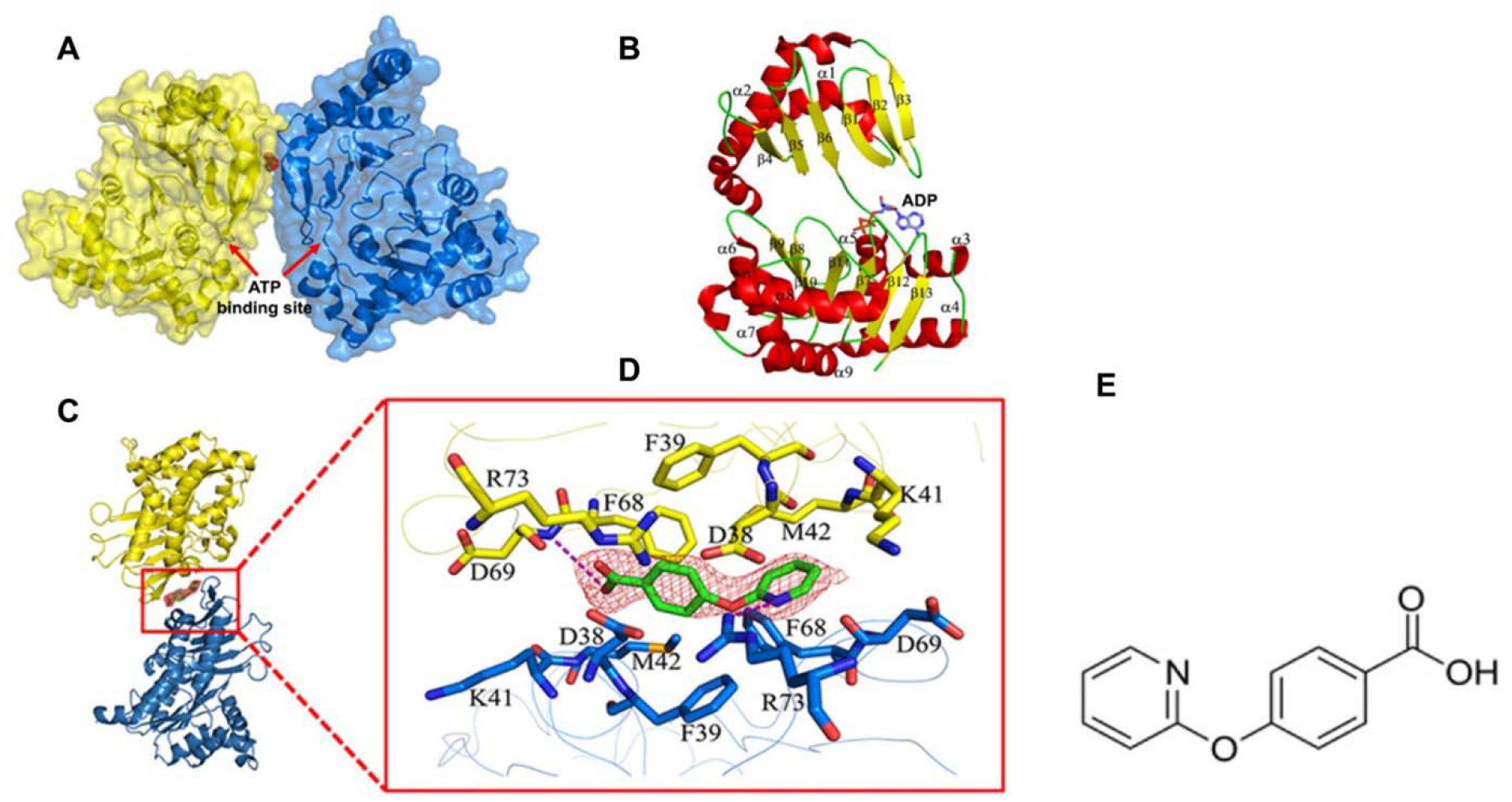
1G2 Binding determined by X-ray crystallography. A) Cartoon representation of the crystal structure of Cagα crystallized as two molecules in the asymmetric unit. Red map in middle of two subunits represents molecule 1G2 and arrows indicate the ATP/ADP binding site. B) Representation of the monomeric subunit of Cagα in ribbon form: α helices, β strands and loops are represented in yellow, red and green, respectively. The nine helices are labeled as α1 to α9 and the β-strands are labeled as β1 to β13 and the ADP binding site is indicated. C) Side view of the interaction of two subunits of protein with 1G2 in the middle represented as green stick and red map. D) Enlarged view of 1G2 binding at the interface between two protein subunits. The 2FO-FC electron density map of 1G2 was contoured at 1.5σ. E) 2D structure of 1G2.

### Validation of the 1G2 binding site by mutagenesis of Cagα

To validate the binding site identified by X-ray crystallography (Fig. 1D) we mutagenized the *cag*α gene to change five amino acids at the binding site. We then purified the binding site mutants CagαD38A, CagαF39A, CagαK41A, CagαM42A, CagαR73A, CagαR73K and analysed their binding to 1G2 using differential scanning fluorimetry (DSF). This technique determines the melting temperature of a protein in a temperature gradient in the presence of the fluorescent dye SYPRO orange. Positive shifts are generally considered as evidence for binding due to stabilization by a small molecule (37). Analysis of the melting temperatures suggested that CagαD38A and CagαM42A bind 1G2 like the wild type protein whereas the binding of Cagα mutants F39A, K41A, R73A and R73K was reduced (Fig. 2A). These results suggest that Cagα residues F39, K41 and R73 contribute to binding of 1G2.

**Figure 2:**
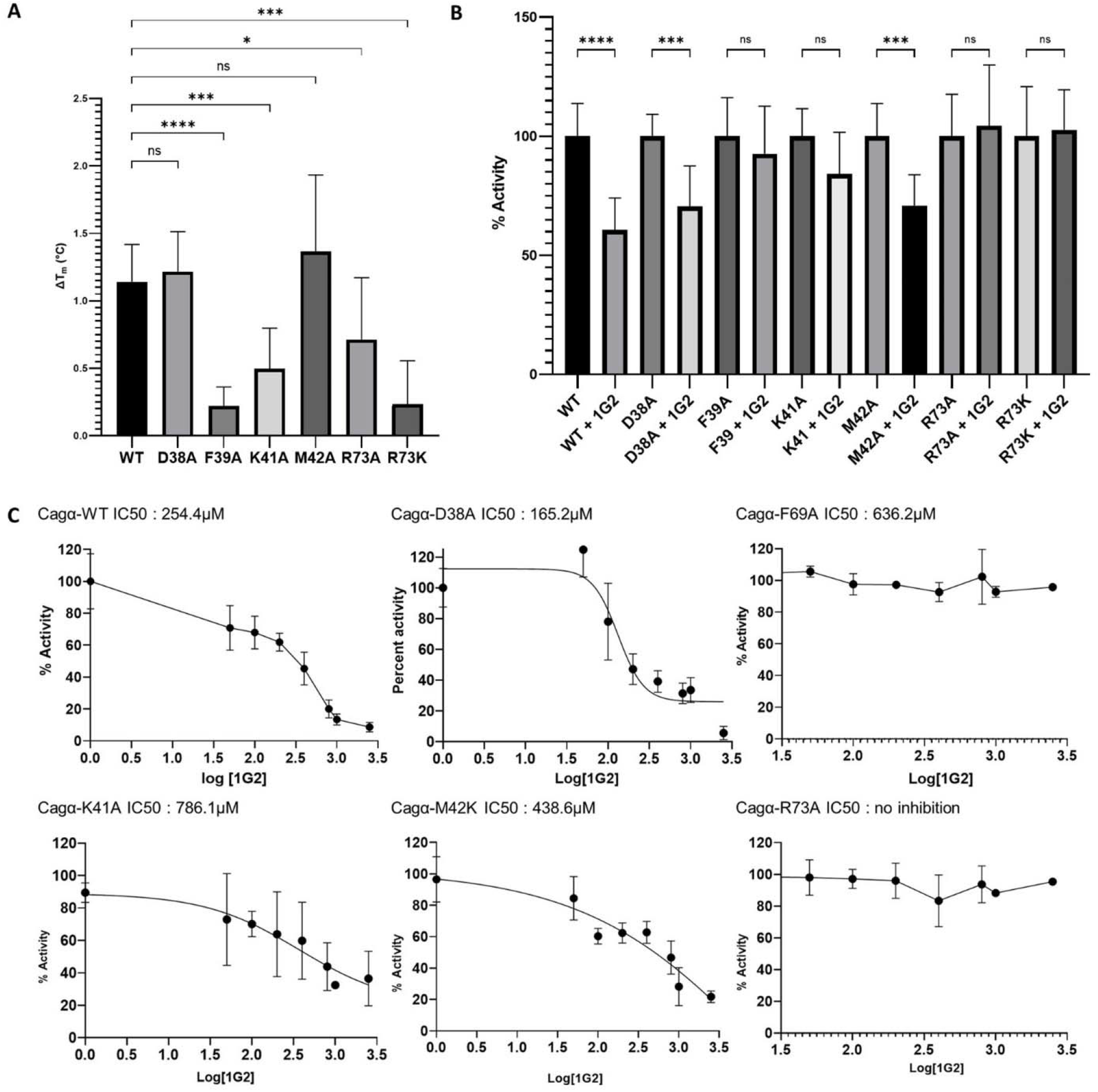
Characterization of active site mutants by differential scanning fluorimetry and ATPase enzyme assays. A) Changes in the melting temperature of Cagα mutant proteins were measured by DSF after incubation with molecules at 5 mM final concentration (n= 3 times triplicates). The results were analysed using ANOVA test ns ; non significant, * p<0.05, *** p=<0.001, **** p<0.0001. B) Effect of 1G2 derivatives on the ATPase activity of Cagα mutant proteins. Cagα WT and mutant proteins were incubated with 1G2 at different concentrations and ATPase activity was measured by malachite green assay, ANOVA test ns ; non significant, *** p=<0.001, **** p<0.0001. C) Dose-response curves of ATPase activity of mutant proteins showing IC_50_ values in the presence of 1G2. C) Dose-response curves of ATPase activity of mutant proteins showing IC_50_ values in the presence of 1G2.

We next determined the ATPase activity of purified Cagα and its mutants using a malachite green assay measuring the release of inorganic phosphate from the ATP. All Cagα mutants have ATPase activities similar to the wild type protein and incubation with 1G2 at a concentration of 200 µM reduced ATPase activity of most of them with the exception of CagαR73A that was not inhibited (Fig. 2B). The effect of 1G2 on CagαF39A, CagαK41A and CagαM42A was less pronounced than on the wild type protein. To gain more quantitative information on the effect of 1G2 on the different mutants we varied the concentration of the inhibitors and calculated IC_50_ values as a measure of inhibitor efficacy. The IC_50_ values for inhibition of the wild type and of CagαD38A are in a similar range, those of CagαK41A and CagαM42A are significantly higher and CagαF39A and CagαR73A are resistant to the inhibitor (Fig. 2C). The enzyme assay results are consistent with those obtained by DSF suggesting that Cagα residues F39 and R73 are critically important for binding of 1G2 and that K41 and M42 also contribute to the effect of the inhibitor.

### Design and selection of 1G2 derivatives molecules

Based on the results from co-crystallisation and mutagenesis of the active site we created a library of 96 1G2 derivatives by medicinal chemistry in an attempt to generate more potent inhibitors. We designed 1G2 derivatives to improve binding to amino acid residues at the inhibitor binding site and we also introduced functional groups that may modulate solubility and thereby permeation into cells (Fig. 3 and suppl. Fig. 3). The fact that a previously identified 1G2 derivative (1G2#4) (37) was more efficient against Cagα *in vitro* also informed this strategy since its additional methyl group may improve binding to K41. As first step to identify more potent 1G2 derivatives we screened for toxicity for human AGS (gastric adenocarcinoma) cells that are used for *H. pylori* infection experiments. Using a cell proliferation assay we identified four cytotoxic molecules, triggering more than 20% of mortality and these molecules were not further investigated. Next, we used a bacterial growth inhibition assay to test toxicity for *H. pylori* using an antibiogram-like technique with 200 μM of 1G2 and derivatives (chloramphenicol as a positive control) on filter plates and we found that none of the 92 derivates is toxic for the bacteria (suppl. Tab. 4).

**Figure 3:**
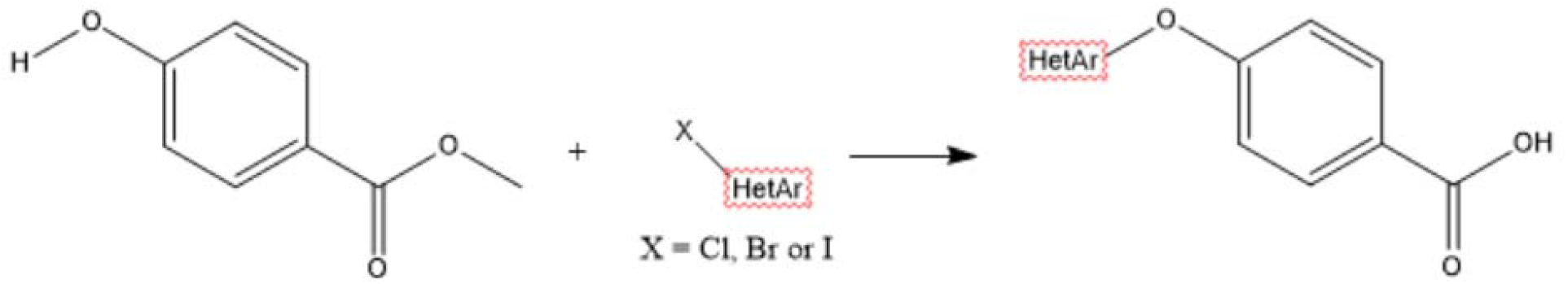
Synthesis of 1G2 derivatives See also suppl. Fig. 3.

To determine the effect of the 1G2 derivatives on T4SS function we quantified the production of interleukin 8 (IL-8) during AGS cell infection by *H. pylori*. IL-8 is an inflammatory factor induced after co-infection with *H. pylori* and it provides a quantifiable readout since it is secreted into the cell culture supernatant (14). We designed a medium-throughput assay enabling us to measure the effects of preincubation of the molecules on IL-8 production of AGS cells in 96-well microtiter plates, followed by quantification using an ELISA assay. We conducted two rounds of screening to identify molecules that reduce IL-8 production more strongly than 1G2 in a reproducible fashion. The first round identified 25 molecules that cause similar or more pronounced reduction of IL-8 production (suppl. Fig. 2A) and a secondary screen identified five molecules (1G2_1313, 1G2_1338, 1G2_2886, 1G2_2889 and 1G2_2902) that lead to strong reductions of IL-8 induction in a reproducible fashion (Tab. 1 and suppl. Fig. 2B). The selected five molecules were further characterized using the infection model described by Arya *et al*. (37) and the assays were repeated six times to gain sufficient statistical power. To compare the effects of the molecules on the T4SS-dependent IL-8 induction, we subtracted the value of the control Δ*cagV* mutant strain, which represents the background on non-dependent T4SS IL-8 induction. We observed that 1G2 reduces IL-8 production to about 20% as compared to the wild type control without inhibitor. The five selected molecules have varying effect on IL-8 production. Molecules 1G2_2886, 1G2_2889 and 1G2_2902 have comparable effects to 1G2, but 1G2_1313 and 1G2_1338 reduce IL-8 production close to the background level of the Δ*cagV* mutant (Fig. 4).

**Figure 4:**
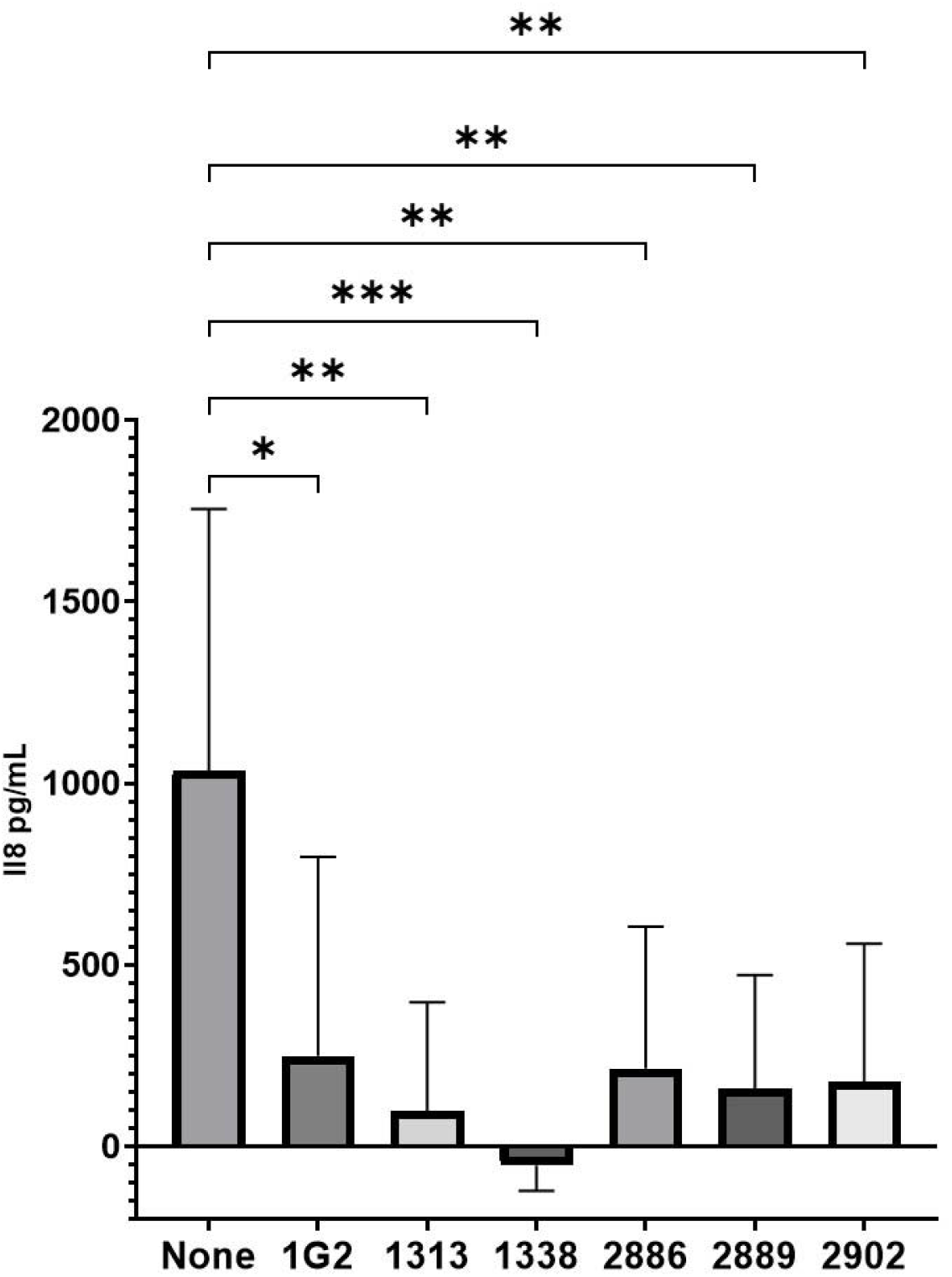
Quantification of the interleukin-8 production by AGS cells infected by *H. pylori* strains after treatment with the 1G2 and its derivatives. AGS cells were infected by the strain 26695 that had been preincubated with 200 μM of molecules for 4 h and the amount of secreted IL-8 was measured by ELISA. The T4SS dependent induction of IL-8 is displayed in pg/ml (n=6) and the value for strain Δ*cagV* were used as baseline. *p<0.01, ** p<0.001, ***p<0.0005, Ordinary one-way ANOVA test

**Table 1:**
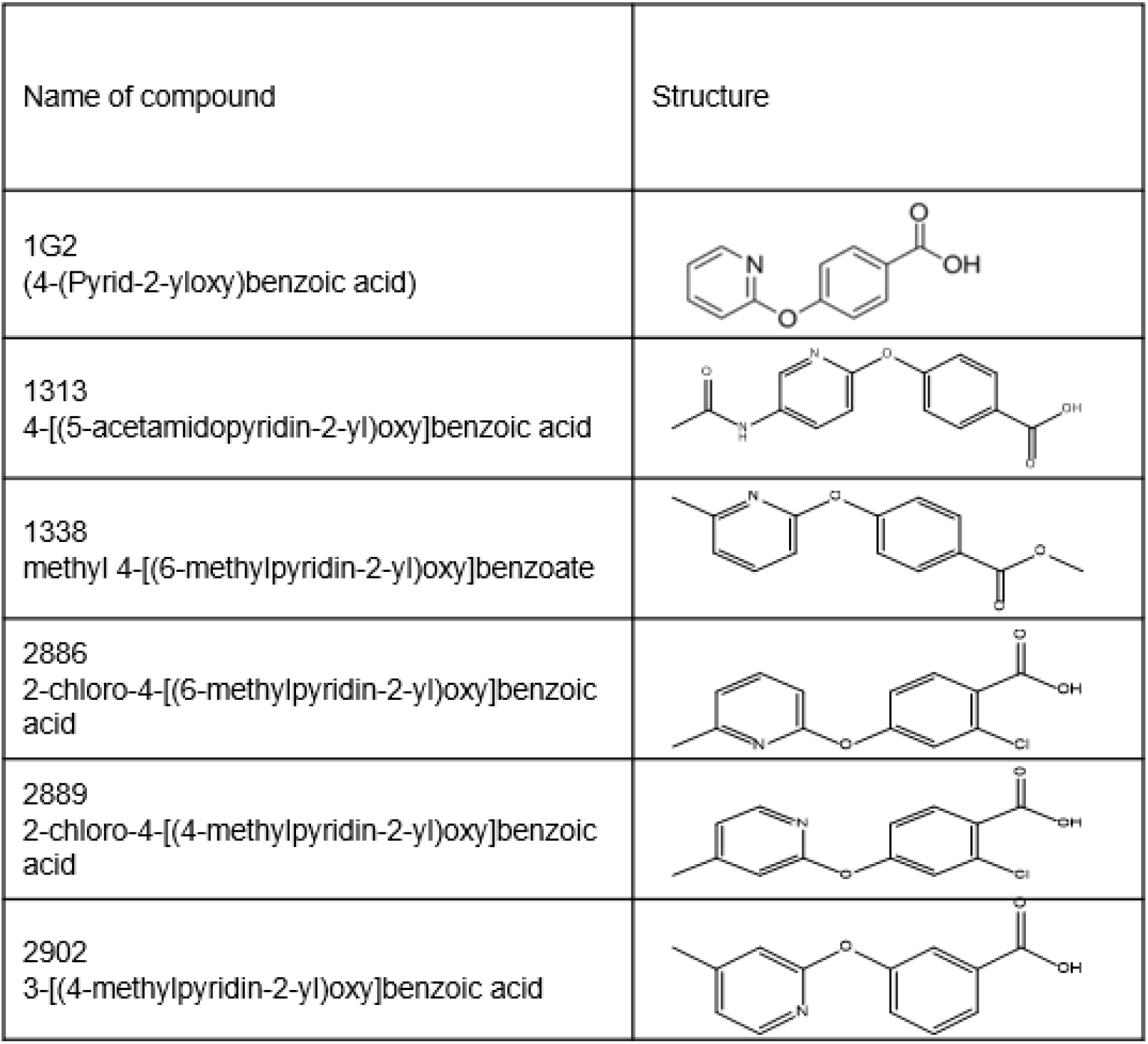
Structures of 1G2 derivates identified by screening for potency in AGS cell infection assays.

| Name of compound | Structure |
| --- | --- |
| 1G2<br>(4-(Pyrid-2-yloxy)benzoic acid) |  |
| 1313<br>4-[(5-acetamidopyridin-2-yl)oxy]benzoic acid |  |
| 1338<br>methyl 4-[(6-methylpyridin-2-yl)oxy]benzoate |  |
| 2886<br>2-chloro-4-[(6-methylpyridin-2-yl)oxy]benzoic acid |  |
| 2889<br>2-chloro-4-[(4-methylpyridin-2-yl)oxy]benzoic acid |  |
| 2902<br>3-[(4-methylpyridin-2-yl)oxy]benzoic acid |  |

### Differential scanning fluorimetry and docking suggest that 1G2 derivatives bind Cagα

To characterize the five selected 1G2 derivatives we first tested their binding to Cagα using the DSF approach as above. Molecules 1G2_1313, 1G2_2889 and 1G2_2902 increased the melting temperature to a lower extent than 1G2, 1G2_2886 had a similar effect as compared to 1G2 and 1G2_1338 actually decreased the melting temperature (Fig. 5). We also conducted *in silico* docking using Autodock Vina software (47). This approach enables a computational prediction of the binding energies and the negative value obtained for 1G2 (-6.6 ΔG/mol) is consistent with its binding to Cagα (suppl. Tab. 3). The predicted binding energies for the five 1G2 derivatives (1G2_1313, 1G2_1338, 1G2_2886, 1G2_2889 and 1G2_2902) had more pronouncedly negative values (-9 to -10.2 ΔG/mol) suggesting that they may have higher affinities for Cagα than 1G2. Whereas the DSF results are not consisted with higher affinity of the 1G2 derivatives to Cagα the docking results suggest stronger binding and we tested their effects on enzyme activity next.

**Figure 5.**
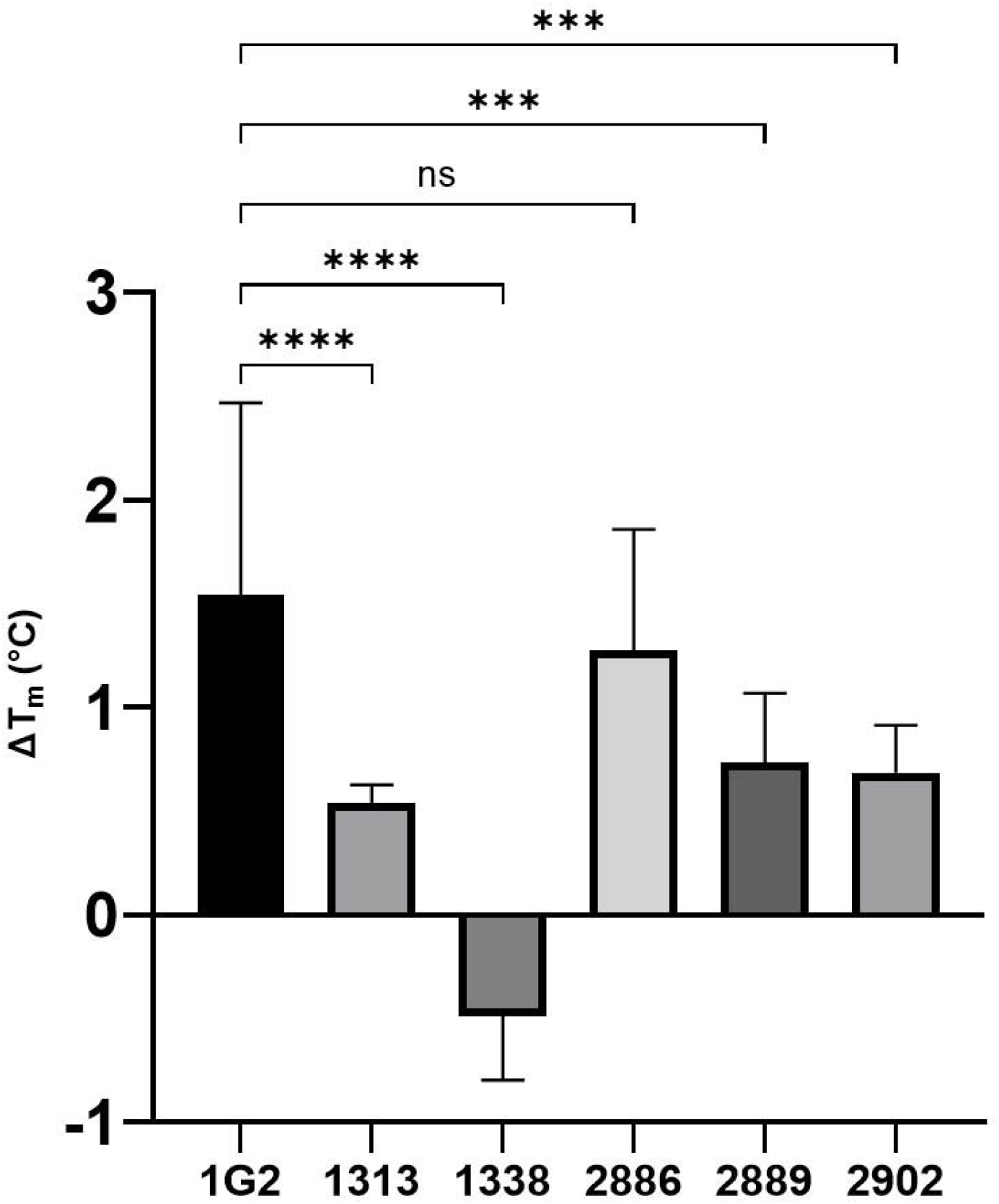
Melting temperature of Cagα in the presence of ligands. Melting temperatures for Cagα were determined using differential scanning fluorimetry (DSF). ANOVA test, ns ; non significant, *** p=<0.001, **** p<0.0001

### Effect of 1G2 derivatives molecules on the ATPase activity of Cagα

Using the malachite green assay as above we assessed to which degree the five 1G2 derivatives impact the enzymatic activity of Cagα. All molecules reduce the Cagα ATPase activity to varying degrees when they were applied at 500 μM concentration (Fig. 6A). We then tested the inhibitors at varying concentrations and the IC_50_ value of molecule 1G2_1313 is similar to that of 1G2 (256 μM). In contrast, molecules 1G2_1338 (60 μM), 1G2_2886 (47.50 μM), 1G2_2889 (96 μM) and 1G2_2902 (85μM) have lower IC_50_□values than 1G2. These results are consistent with the increased negative effects of these derivatives observed in the IL-8 production assay (Fig. 4) and further support the notion that these derivatives are more potent than 1G2.

**Figure 6.**
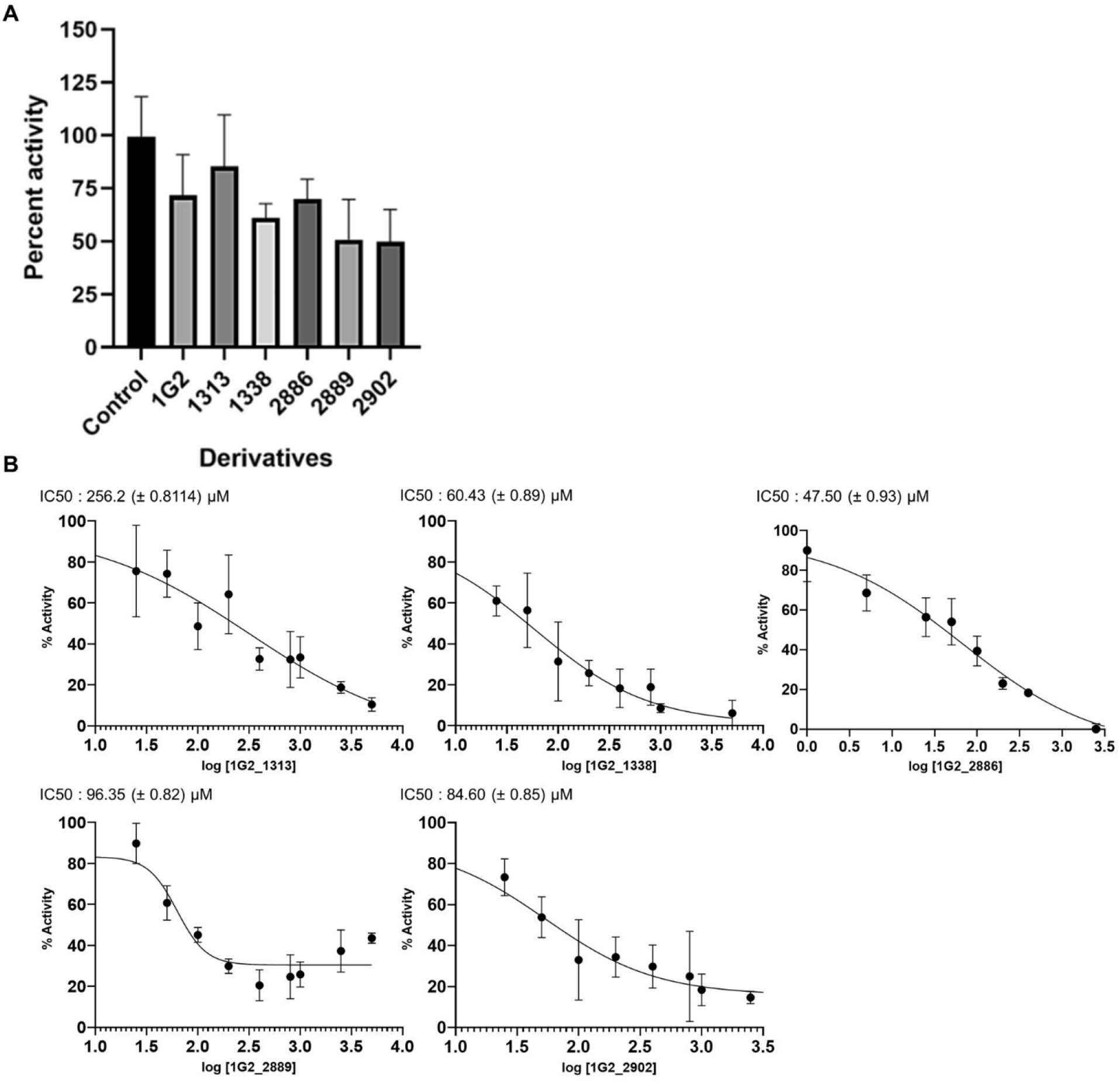
Enzyme assays of Cagα in the presence of molecule 1G2 and derivatives. A) Inhibition of ATPase activity of Cagα in the presence of 500 μM concentration of molecules, representation of one triplicate assay. B) Dose-response curves of ATPase activity showing IC_50_ values in the presence of 1G2 and its derivatives 1G2_1313, 1G2_1338, 1G2_2886, 1G2_2889 and 1G2_2902.

### 1G2 and derivatives affect the assembly of T4SS pili

Finally, we tested the effects of 1G2 and its five derivatives on the physiology of the T4SS. AGS cells were infected with *H. pylori* wild type strain 26695 that had been pre-incubated with 200 μM of the molecules, followed by infection of 4 hours and analysis of the cells. First, the presence of Cag proteins was analysed using western blotting to determine whether 1G2 and its derivatives have negative impacts on T4SS expression or stability. We do not observe any significant and reproducible effects on the amounts of the translocated virulence factor CagA or of the target Cagα suggesting that the inhibitors do not destabilize the T4SS (suppl. Fig. 4). Second, we used a recently developed quantifiable approach to analyze the T4SS-dependent assembly of extracellular pili using scanning electron microscopy (SEM) during AGS cell infection (Oudouhou and Morin, *et al*, 2023 submitted) (Fig. 7A). We observe on average of 17 (±8) T4SS pili on the wild type strain in the absence of 1G2 and very few on the T4SS-defective Δ*cagV* strain 1 (±2)(Fig. 7B). In contrast, in the presence of 1G2 we observe on average 3(±3) T4SS-pili on the wild type strain and all 1G2 derivatives have similar strong inhibitory effects on pilus assembly (Fig. 7B). These results confirm that 1G2 and its derivatives have strong negative effects on T4SS pilus assembly, and it will be interesting to gain more detailed insights into the molecular mechanism of these molecules in future.

**Figure 7:**
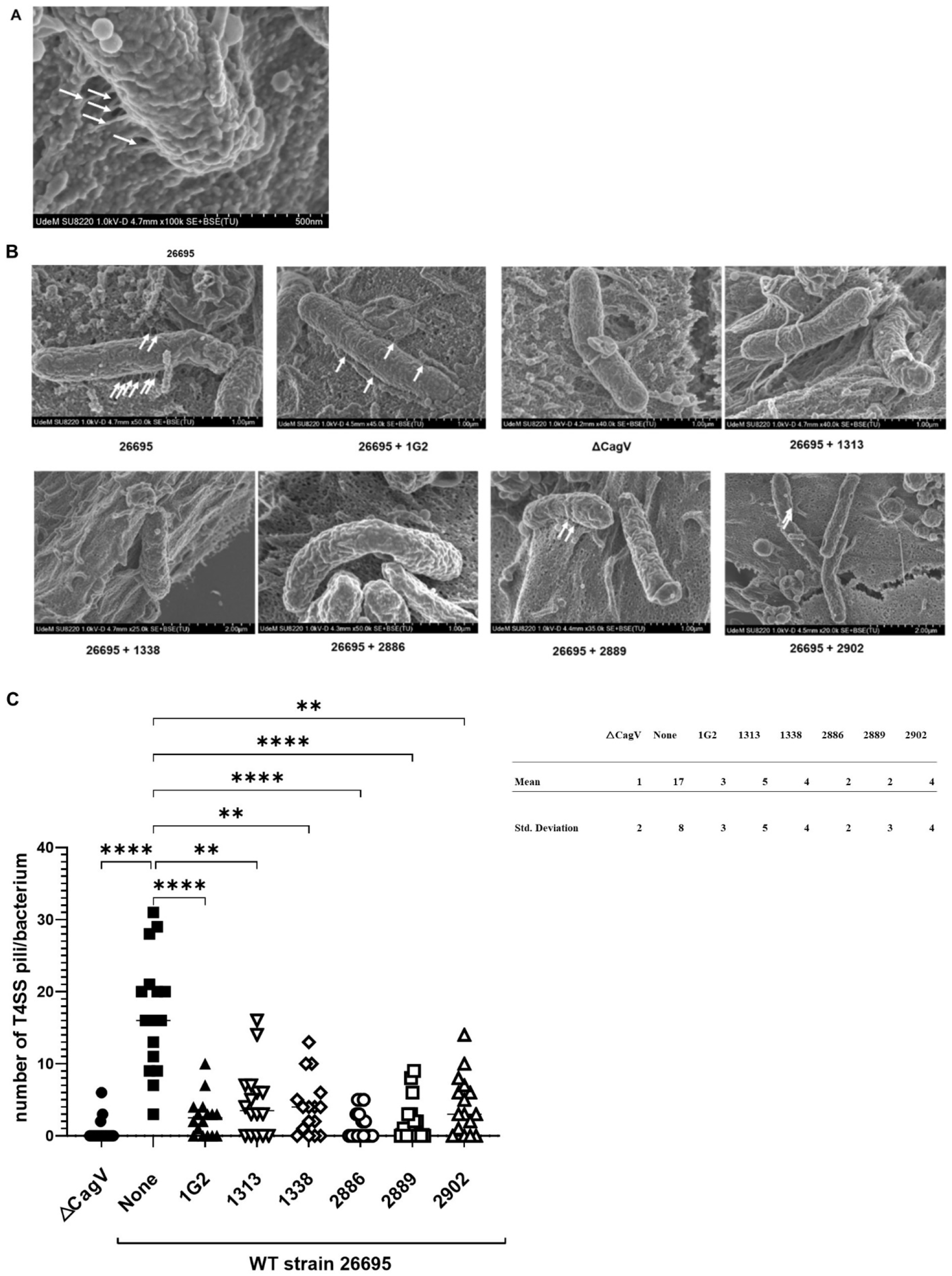
1G2 and its derivates affect the number of T4SSpili during gastric cell infection. AGS cells were preincubated with 1G2 and derivatives and infected with *H. pylori*. A) and B) The T4SSpili were observed using scanning electron microscopy; the T4SS pili are identified with white arrows. C) The T4SS pili were counted using ImageJ. The Δ*cagV* strain was used as negative control. (n=16 bacteria from 2 different infections). ****: p<0.0001,. **p<0.05, ns: non-significant, Kruskal-Wallis ANOVA test

## Discussion

The eradication therapies of *H. pylori* using antibiotics induce a high selection pressure leading to the acquisition of antibiotic resistance that can be transmitted between different strains and other members of the microbiome (24,48–51). Targeting the nonessential virulence factors that the bacteria use to infect the host is an alternative approach to reduce the pressure of selection induced by antibiotics (52–56). In this study we have characterized the binding site of Cagα inhibitor 1G2 using X-ray crystallography and mutagenesis suggesting a unique mechanism of inhibition via a binding site that is distinct from the ATPase active site. X-ray analysis suggested that five amino acids (D38, F39, K41, M42, R73) may form the 1G2 binding site. We mutated them in individually and DSF as well as ATPase enzyme assays showed that F39 and R73 are critically important, and that K41 and M42 also contribute to the binding of 1G2. These data suggest that we have identified the 1G2 binding site and that it functions as an inhibitor of ATPase activity by not directly interfering with substrate binding.

Based on the information gained on the inhibitor binding site we synthesized a series of 98 1G2 derivatives that may have improved binding to amino acids at the binding site and/or penetration into the cells. We tested their toxicity for mammalian cells first to focus our efforts on molecules that do not interfere with other metabolic pathways in mammalian cells (57). In the spirit of our overall anti-virulence strategy, we also validated that these molecules are not toxic for *H. pylori.* We then screened the effects of 1G2 derivatives in a cell-based infection assay monitoring IL-8 production as a readout of T4SS activity to identify more potent anti-virulence molecules. Using this approach, we aimed at identifying 1G2 derivatives that have increased potency for the inhibition of Cagα and/or improved capacity to penetrate into the cells. This strategy was informed by previously identified derivative 1G2#4 that was more active as an inhibitor of Cagα ATPase activity *in vitro* than 1G2 (IC_50_ of 82 μM as compared to 196 μM), but the molecule was inactive as an inhibitor of IL-8 production (37), likely due to solubility issues. This molecule carries an additional methyl group that may form additional hydrophobic interactions with the carbon chain of K41. Using screening of IL-8 production instead of inhibition of ATPase activity as a readout we avoided the identification of molecules like 1G2#4 that are more potent enzyme inhibitors, but inactive as inhibitors of the pathogen.

We then characterized the *in vitro* activity of the five molecules that have the strongest negative effects on IL-8 production as compared to 1G2 (1G2_1313, 1G2_1338, 1G2_2886, 1G2_2889 and 1G2_2902). Interestingly, when we used DSF as an estimate for binding to the target Cagα, none of the molecules led to stronger positive shifts of the melting temperature than 1G2. This suggests that the improved inhibitory activity in the IL-8 production assay is not necessarily due to stronger binding to the target. Interestingly, molecule 1G2_1338 decreased the melting temperature in the DSF assay suggesting that it may have a distinct effect on its target than 1G2. Analysis of the effects of the 1G2 derivatives on Cagα ATPase activity revealed that four of them are in fact more potent inhibitors than 1G2 (1G2_1338: 60 μM, 1G2_2886: 47 μM, 1G2_2889: 96 μM, 1G2_2902: 85 μM). These values are in the range of what we observed previously for 1G2#4, and in addition these molecules are potent inhibitors of IL-8 production reducing this readout of T4SS activity to baseline level of a Cag protein deletion mutant in the case of 1G2_1338. In addition, like 1G2, the five derivatives strongly reduce the production of extracellular T4SS pili suggesting that they share the same overall mechanism of inhibition.

The molecules identified here are the result of a first round of synthesis in the context of a structure-activity relationship (SAR) study and some are already more potent than 1G2 as enzyme inhibitors or in the IL-8 inhibition assay. The results of this work will inform future rounds of optimization by medicinal chemistry to synthesize molecules that have potential for development into drugs. In addition, since these molecules are inhibitors of the assembly of extracellular T4SS pili they could be applied for mechanistic studies of the T4SS assembly and translocation process. Small molecule inhibitors could be applied to block T4SS at different stages of the infection process enabling us to determine the time at which pilus assembly and effector translocation from *H. pylori* to mammalian cells are required.

## Acknowledgements

This work was supported by grants from the Cancer Research Society, the Charles Bowers Memorial Fund, the Bergeron-Jetté Foundation (CRS, #23404 and #25102) and the Natural Sciences and Engineering Research Council (NSERC, #RGPIN-2017-05123) to C.B. We are grateful to Dr. Dainelys Guadarrama Bell and other members of Dr. Antonio Nanci’s laboratory at the Université de Montréal electron microscopy facility for technical support and assistance. Synchrotron X-ray data were collected at the Cornell High Energy Synchrotron Source (CHESS, MacCHESS beamline F1).

## Supplementary information

**supplementary Figure 1.**
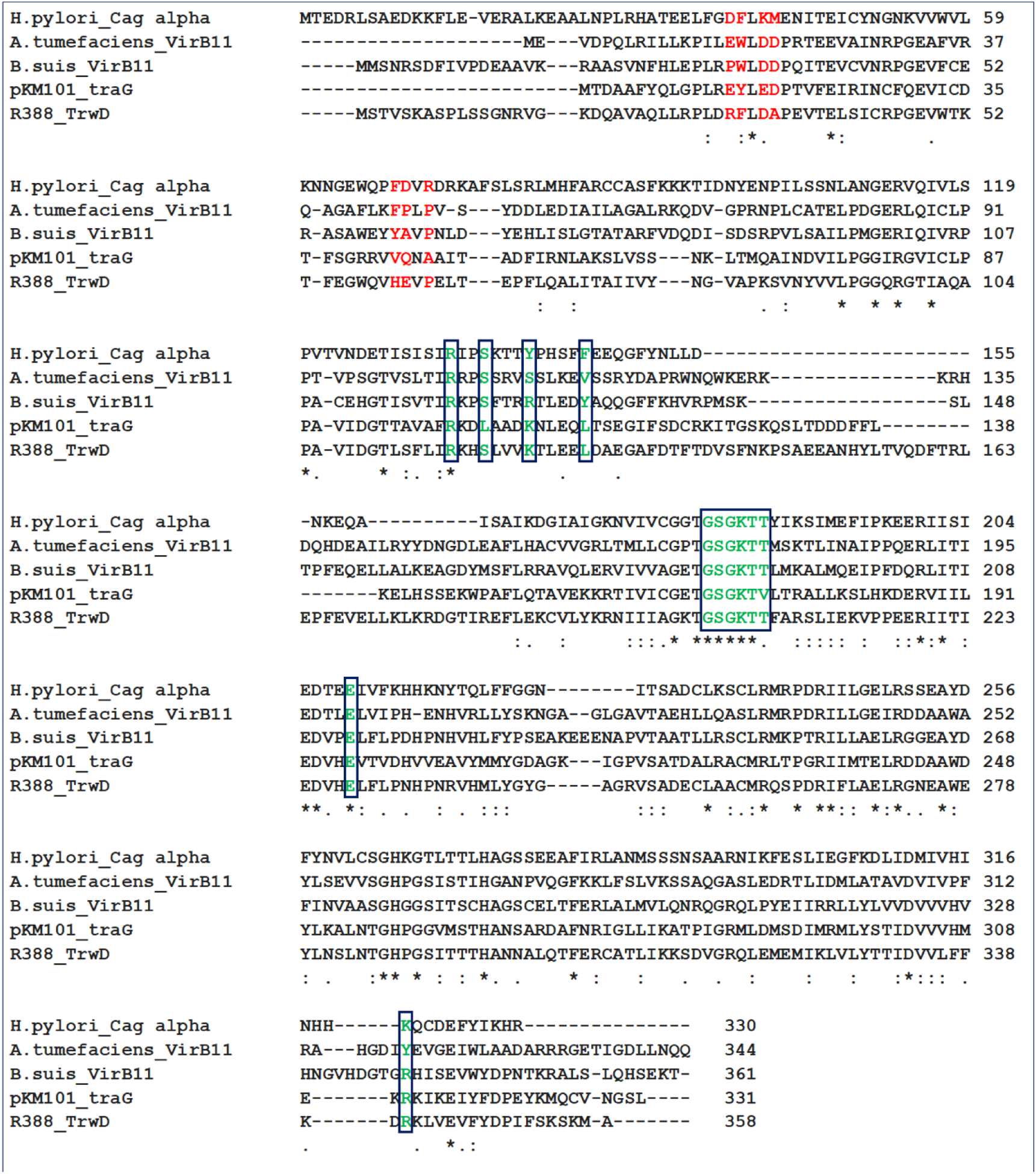
Multiple sequence alignment of Cagα/VirB11 homologs. Selected Cagα/VirB11 homologs from the T4SS of *Agrobacterium tumefaciens* C58, *Brucella suis* 1330, plasmids pKM101 and R388 were aligned using the clustal omega online tool (https://www.ebi.ac.uk/Tools/msa/clustalo/). The amino acids corresponding to the 1G2 binding site on Cagα are indicated in red and the ATPase active site is labelled green with a box.

**supplementary Figure 2:**
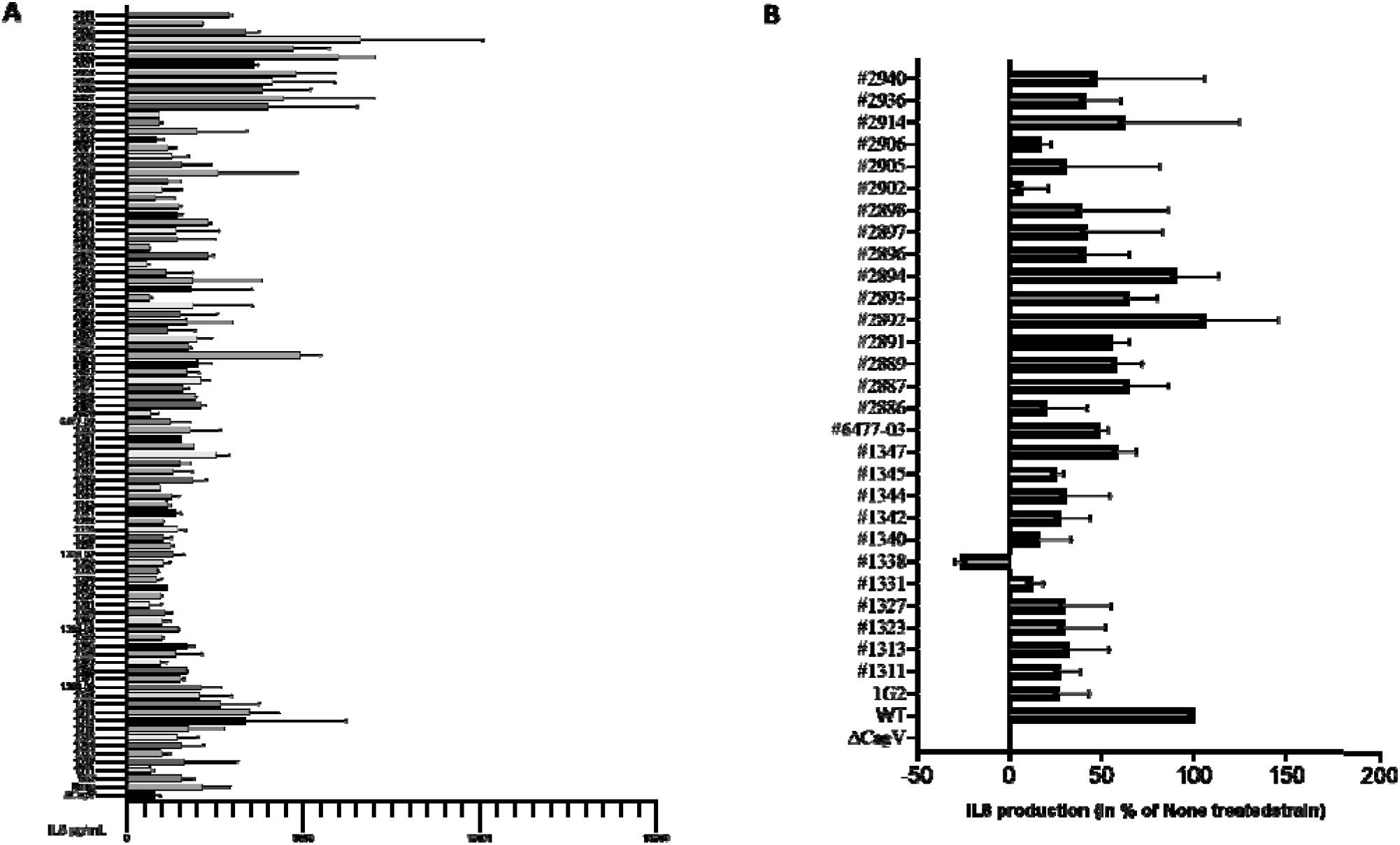
Results of the screens of 1G2 derivatives. AGS cells were infected for 24h with *H. pylori* pre-incubated with 200 μM of molecules and the amounts of secreted IL-8 were measured by ELISA A) first screening results in pg/mL, B) The results of the IL-8 quantification were analyzed by pairs comparing the non-treated result (100% induction) with the treated results. The Δ*cagV* strain was used as negative control and analyzed using the strain 26695.

**supplementary Figure 3:**
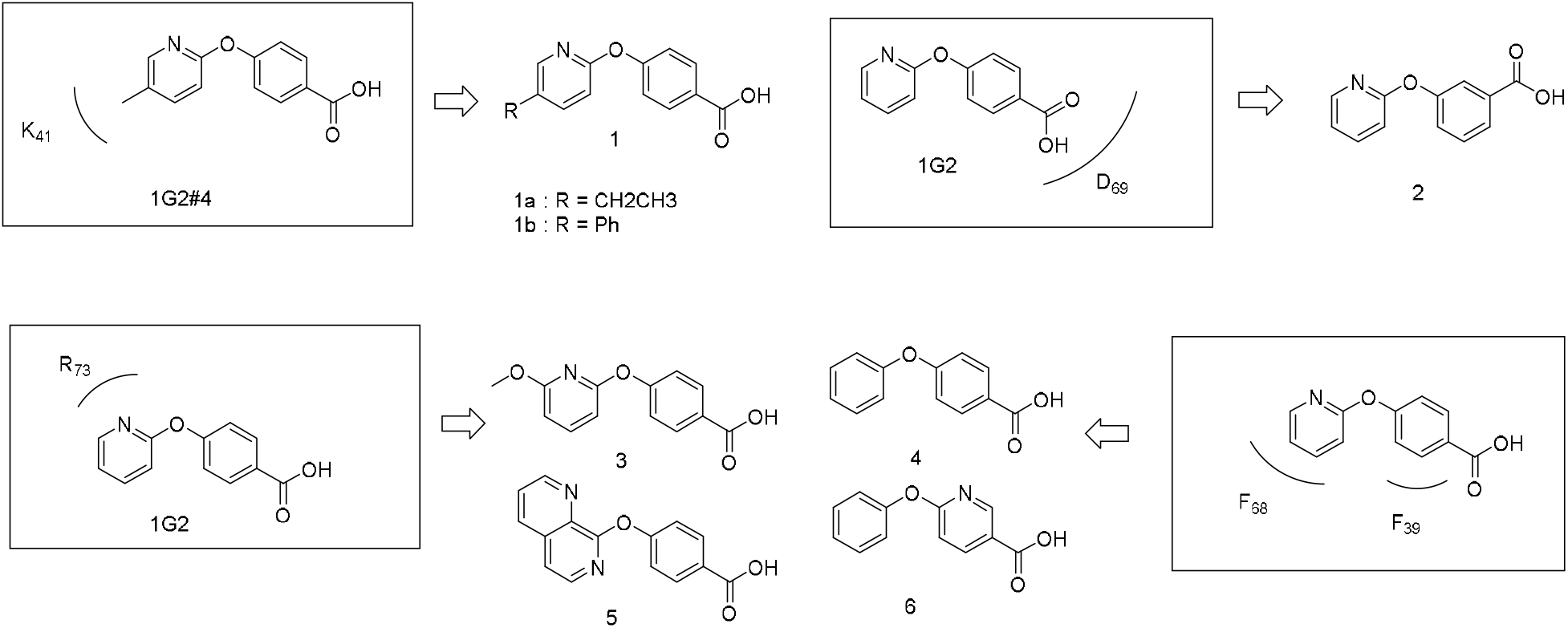
Design of 1G2 analogs. Overview of the synthetic analogs designed to probe the putative interactions between the 1G2 derivates and the various amino acids at the binding site.

**supplementary Figure 4:**
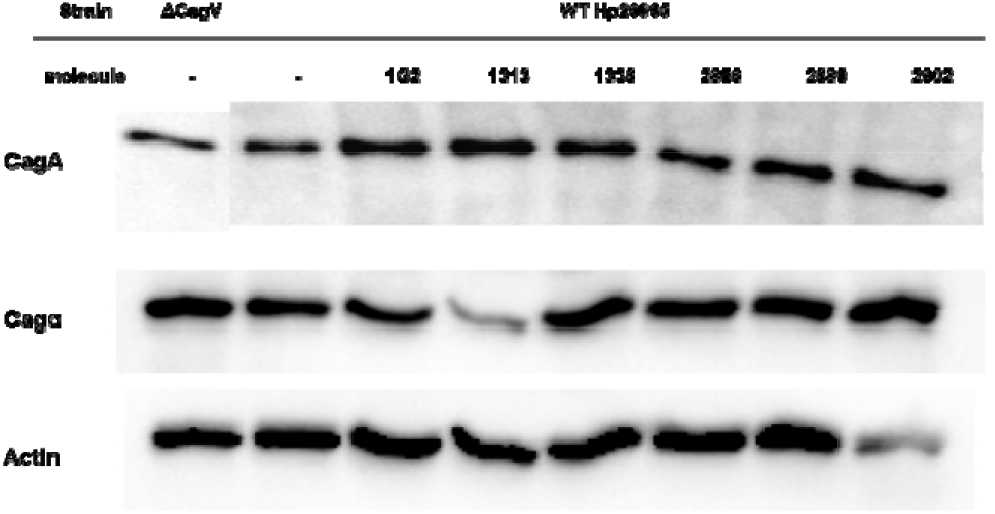
Expression of Cag protein during AGS cells infection with or without treatment with 1G2 and its derivatives. Cell lysates of *H. pylori* strains were separated by SDS-PAGE, followed by western blotting with specific antisera. CagA and Cagα proteins were detected using specific antibodies and actin was used as a loading control. Representative results from four repetitions are shown.

**supplementary Table 1:**
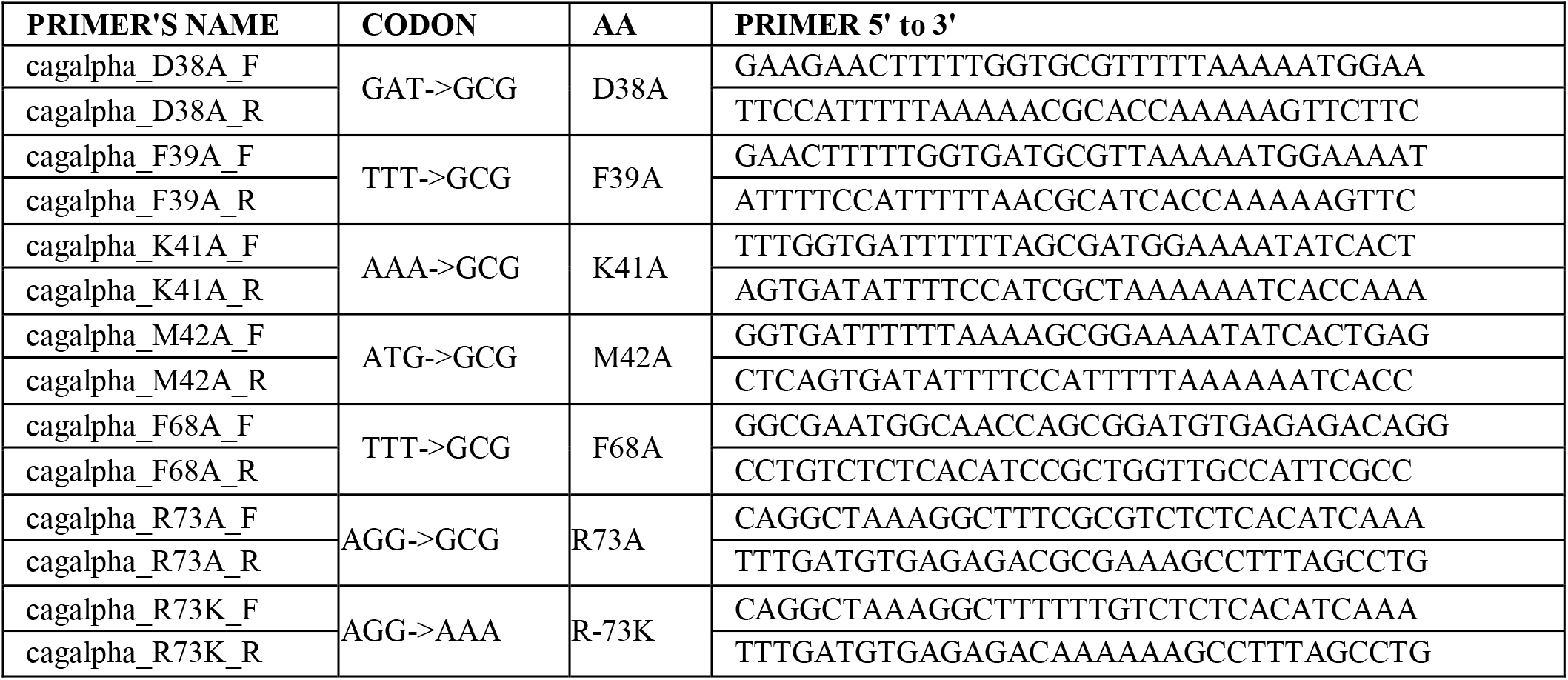
Primers for the creation of 1G2 binding site mutants.

**supplementary Table 2:**
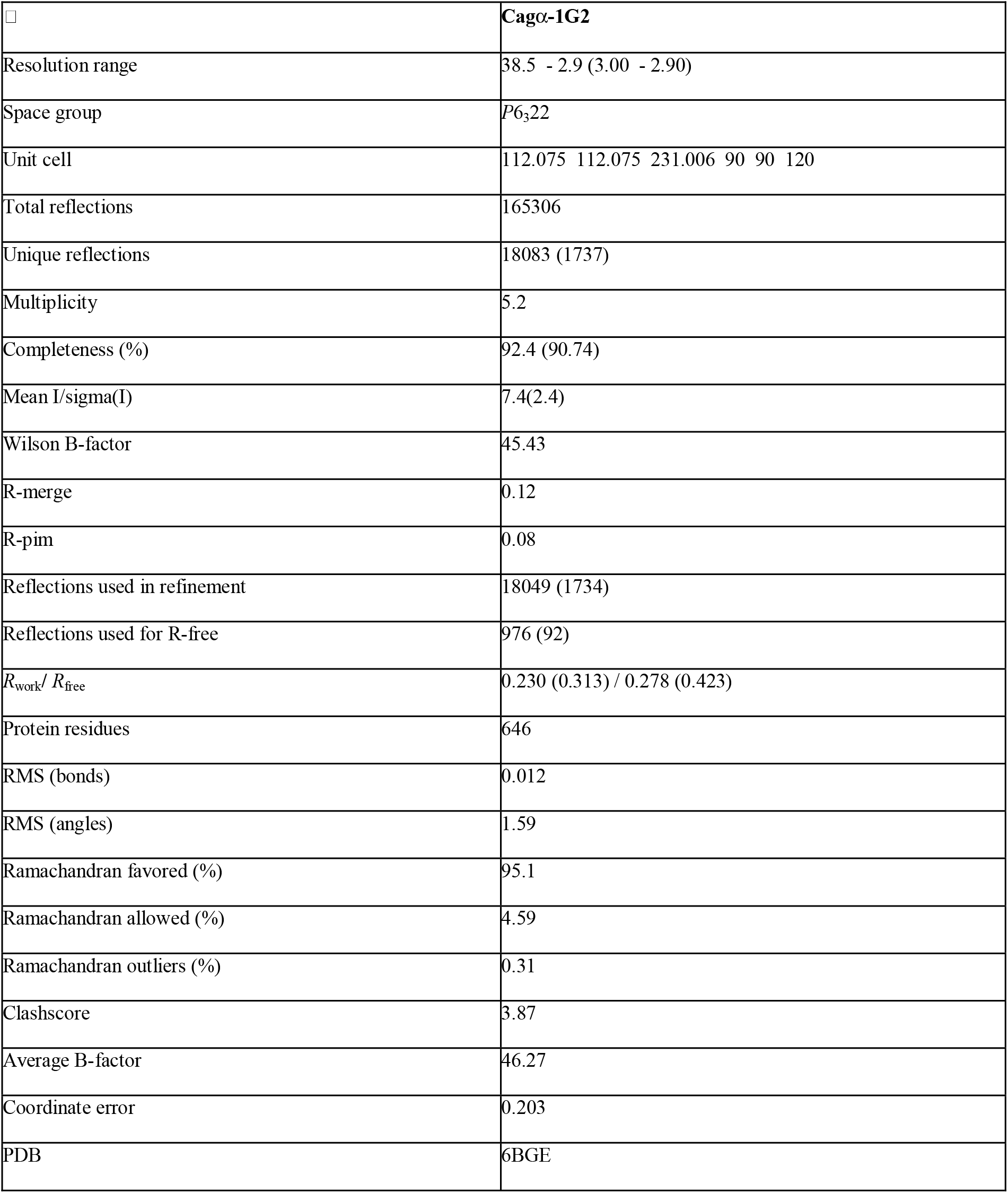
X-ray data collection and refinement statistics.

|  |  |
| --- | --- |
|  | **Cagα-1G2** |
| Resolution range | 38.5 - 2.9 (3.00 - 2.90) |
| Space group | *P*6<sub>3</sub>22 |
| Unit cell | 112.075 112.075 231.006 90 90 120 |
| Total reflections | 165306 |
| Unique reflections | 18083 (1737) |
| Multiplicity | 5.2 |
| Completeness (%) | 92.4 (90.74) |
| Mean I/sigma(I) | 7.4(2.4) |
| Wilson B-factor | 45.43 |
| R-merge | 0.12 |
| R-pim | 0.08 |
| Reflections used in refinement | 18049 (1734) |
| Reflections used for R-free | 976 (92) |
| $R_{\text{work}} / R_{\text{free}}$ | 0.230 (0.313) / 0.278 (0.423) |
| Protein residues | 646 |
| RMS (bonds) | 0.012 |
| RMS (angles) | 1.59 |
| Ramachandran favored (%) | 95.1 |
| Ramachandran allowed (%) | 4.59 |
| Ramachandran outliers (%) | 0.31 |
| Clashscore | 3.87 |
| Average B-factor | 46.27 |
| Coordinate error | 0.203 |
| PDB | 6BGE |

**supplementary Table 3:** Docking of derivatives of molecule 1G2 with target Cagα. Binding energy is shown as kmol/mol as calculated by Autodock Vina software.

| **Molecule** | **Binding energy ($\Delta G/mol$)** |
| --- | --- |
| **1G2** | -6.6 |
| **1G2_1313** | -10.2 |
| **1G2_1338** | -9.0 |
| **1G2_2886** | -9.4 |
| **1G2_2889** | -9.4 |
| **1G2_2902** | -9.6 |

**supplementary Table 4:**
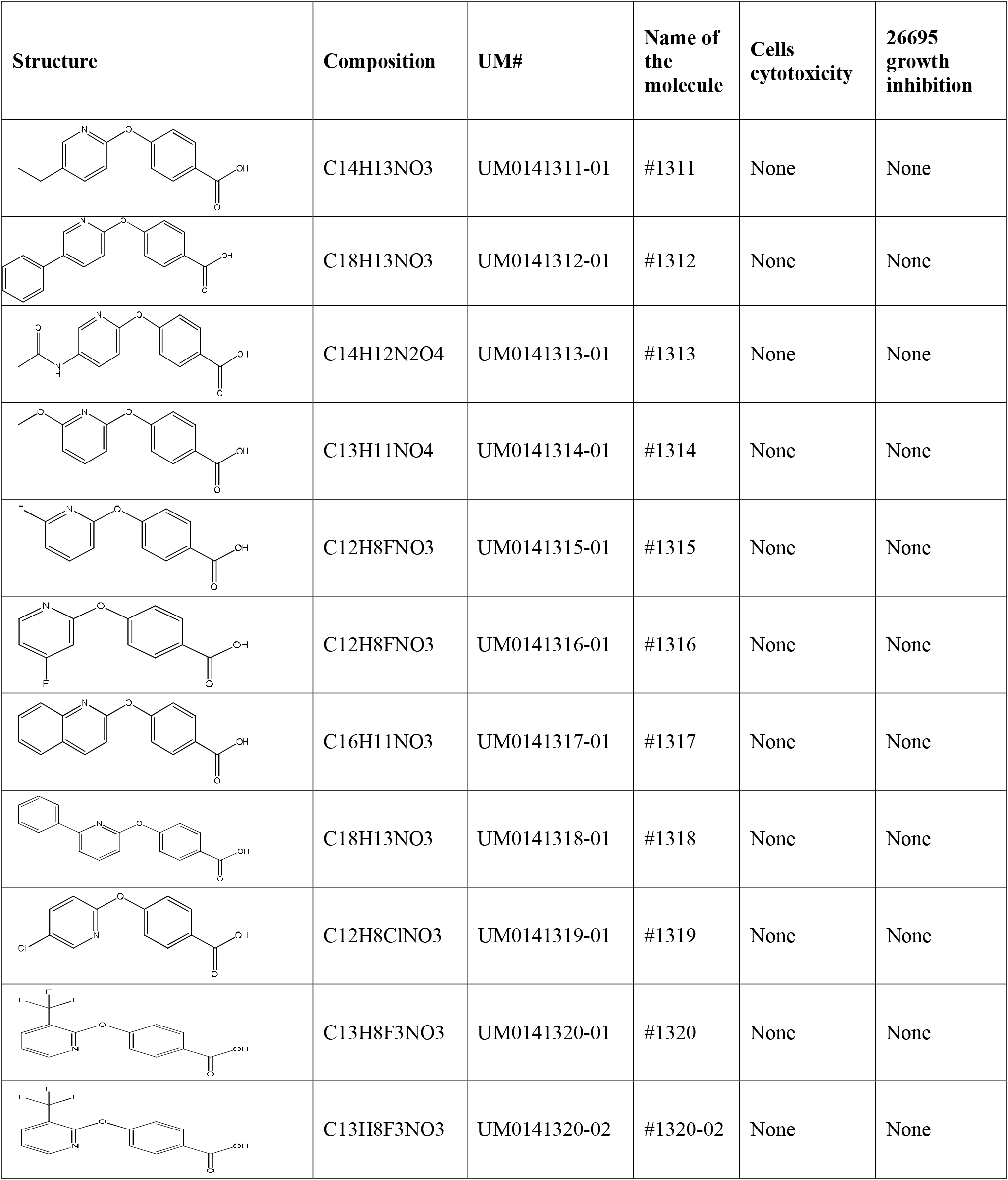
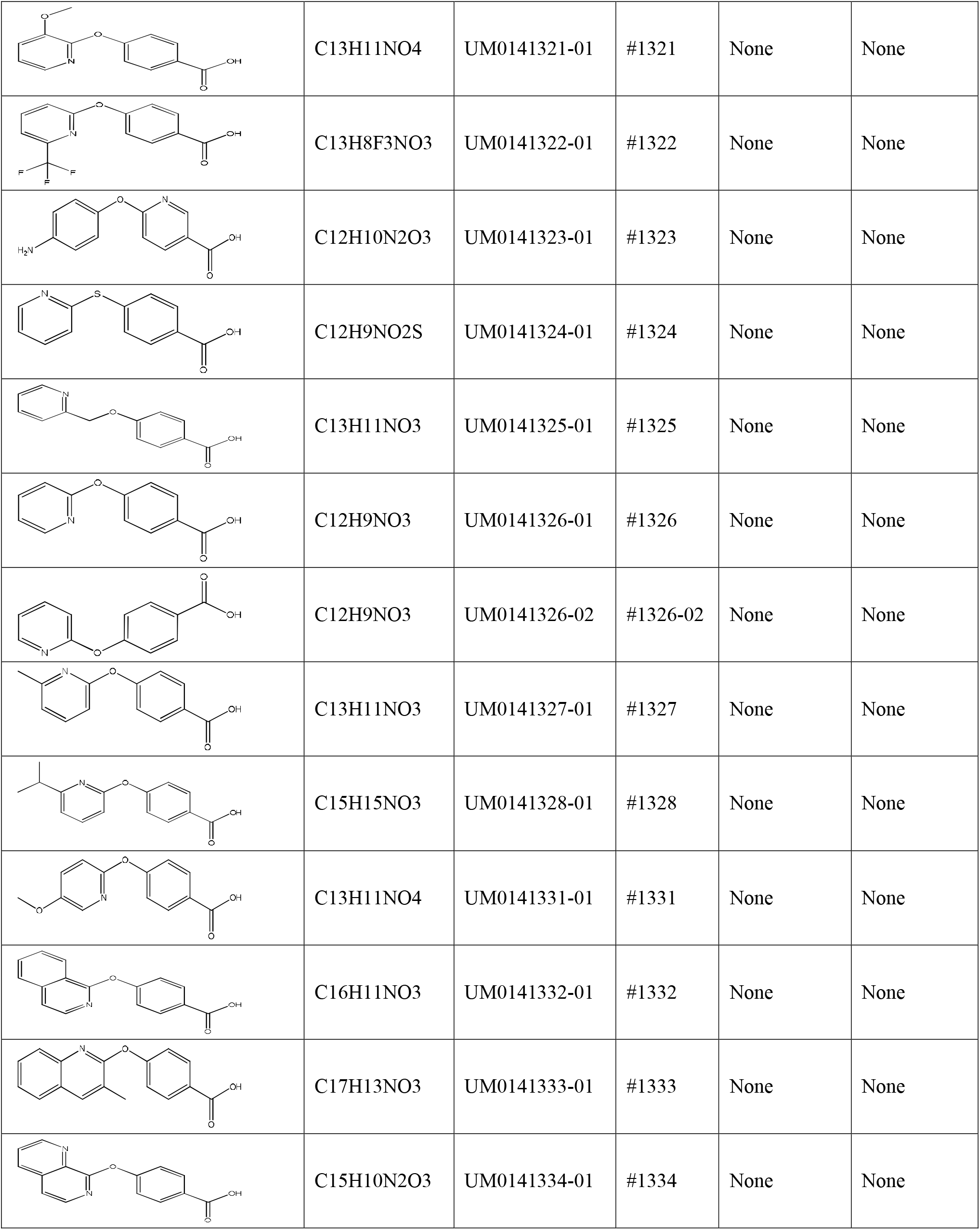
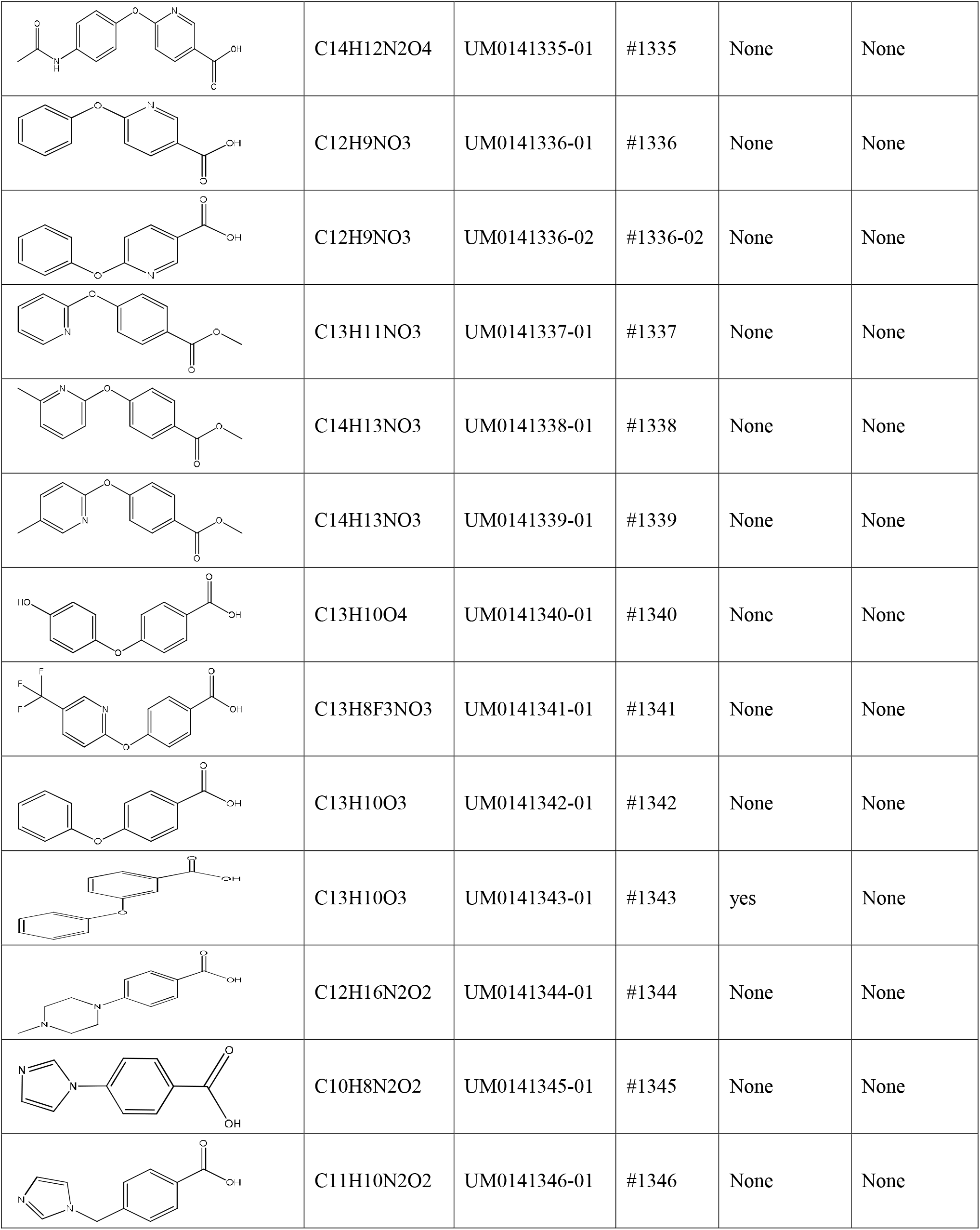
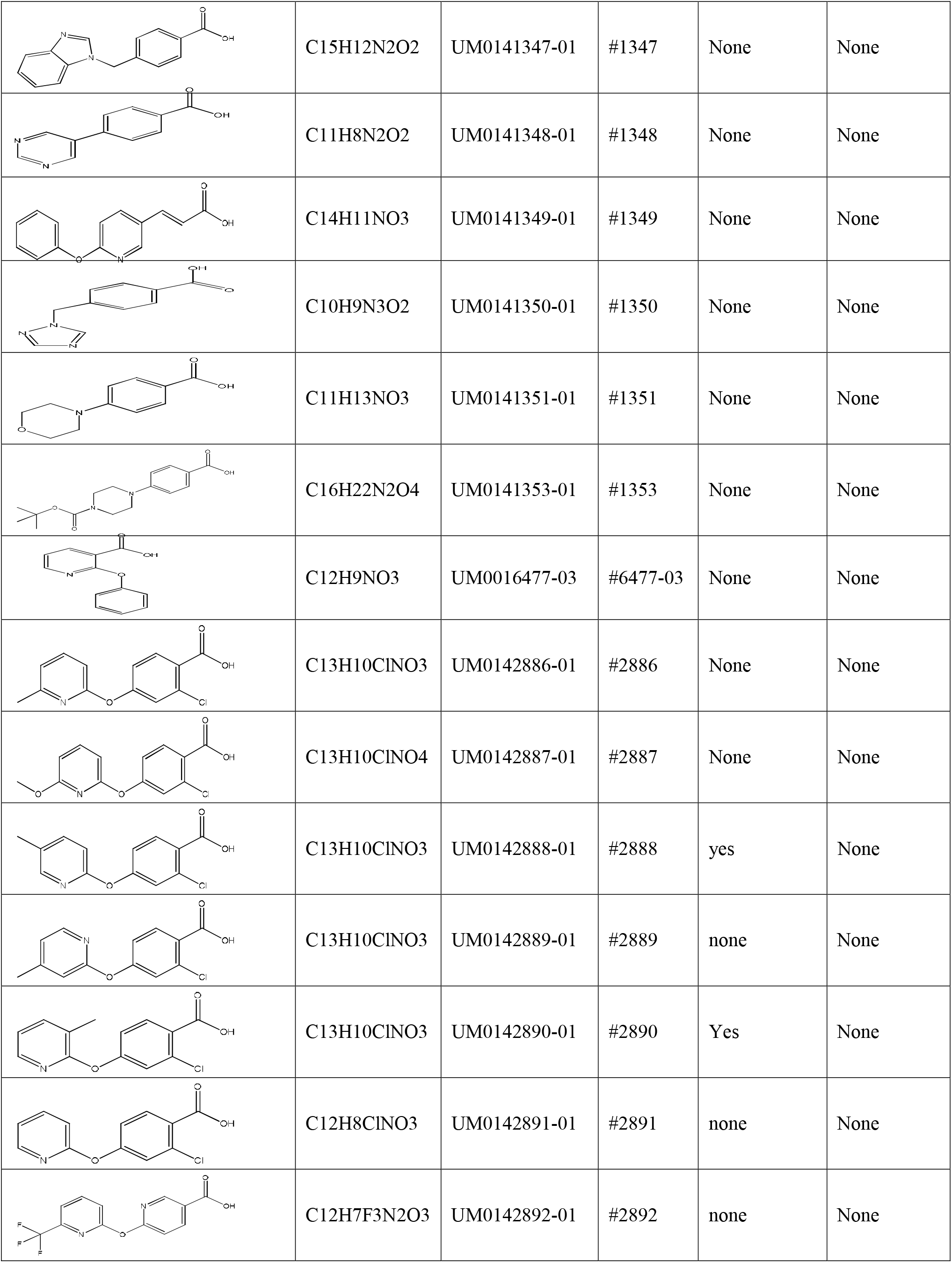
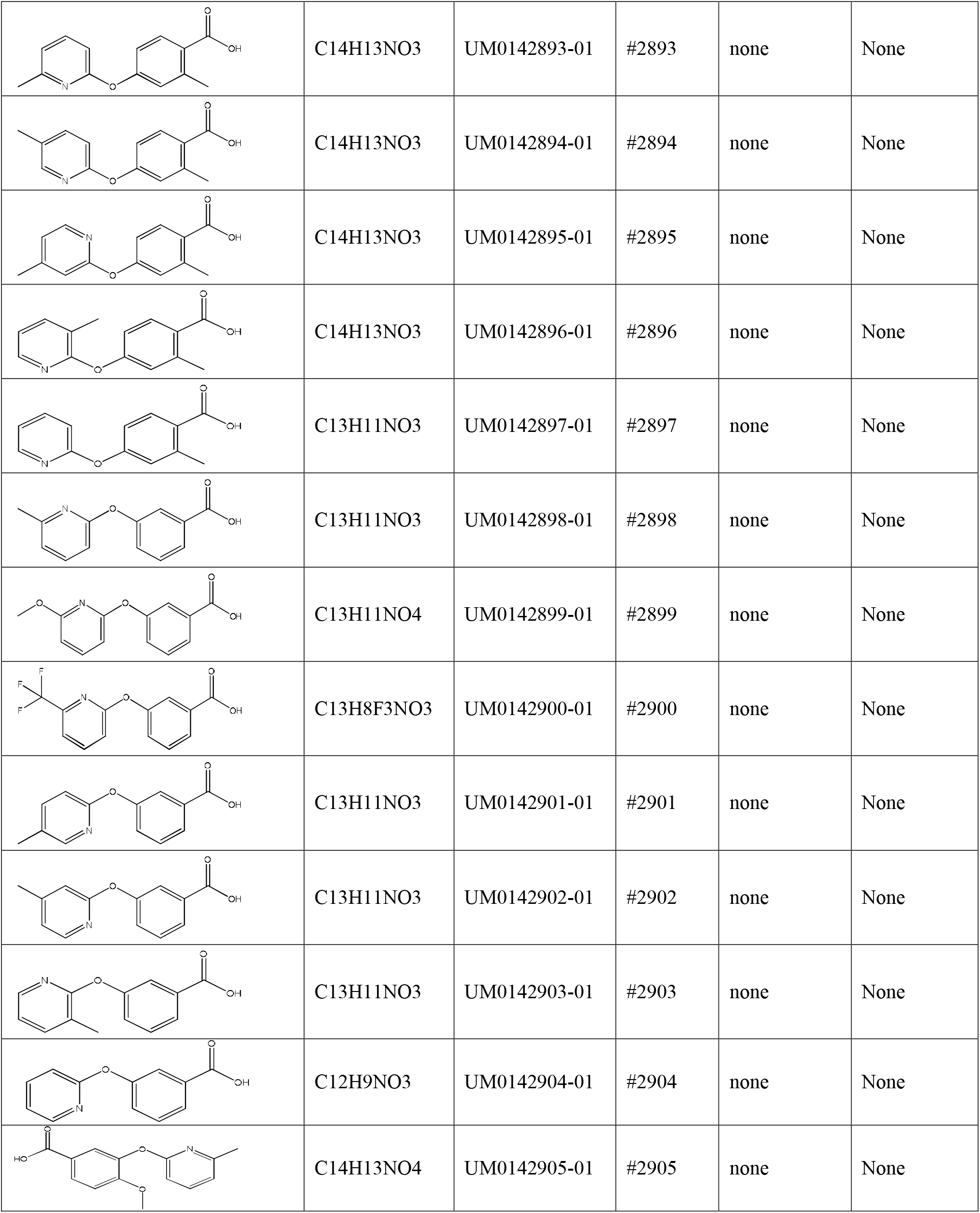
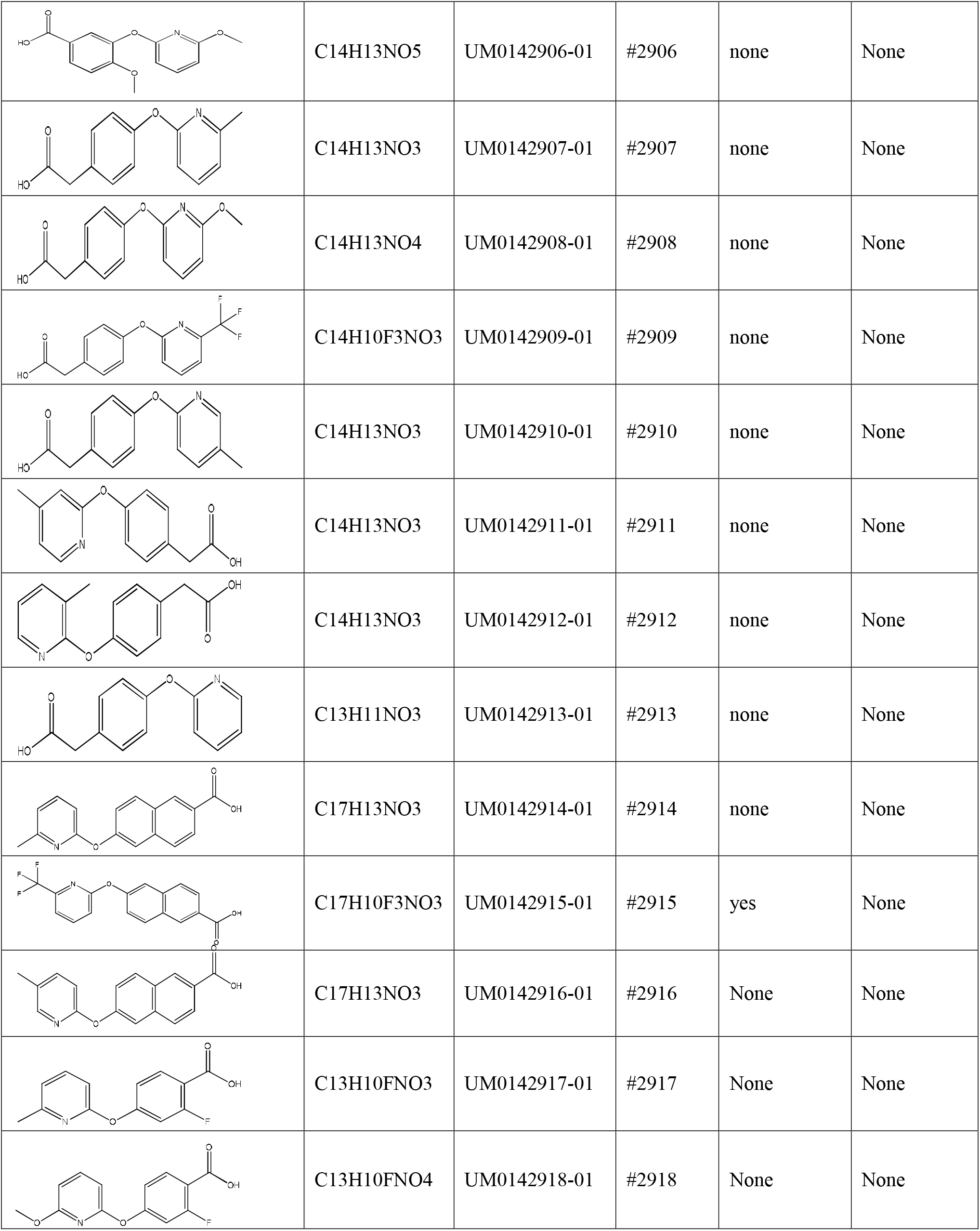
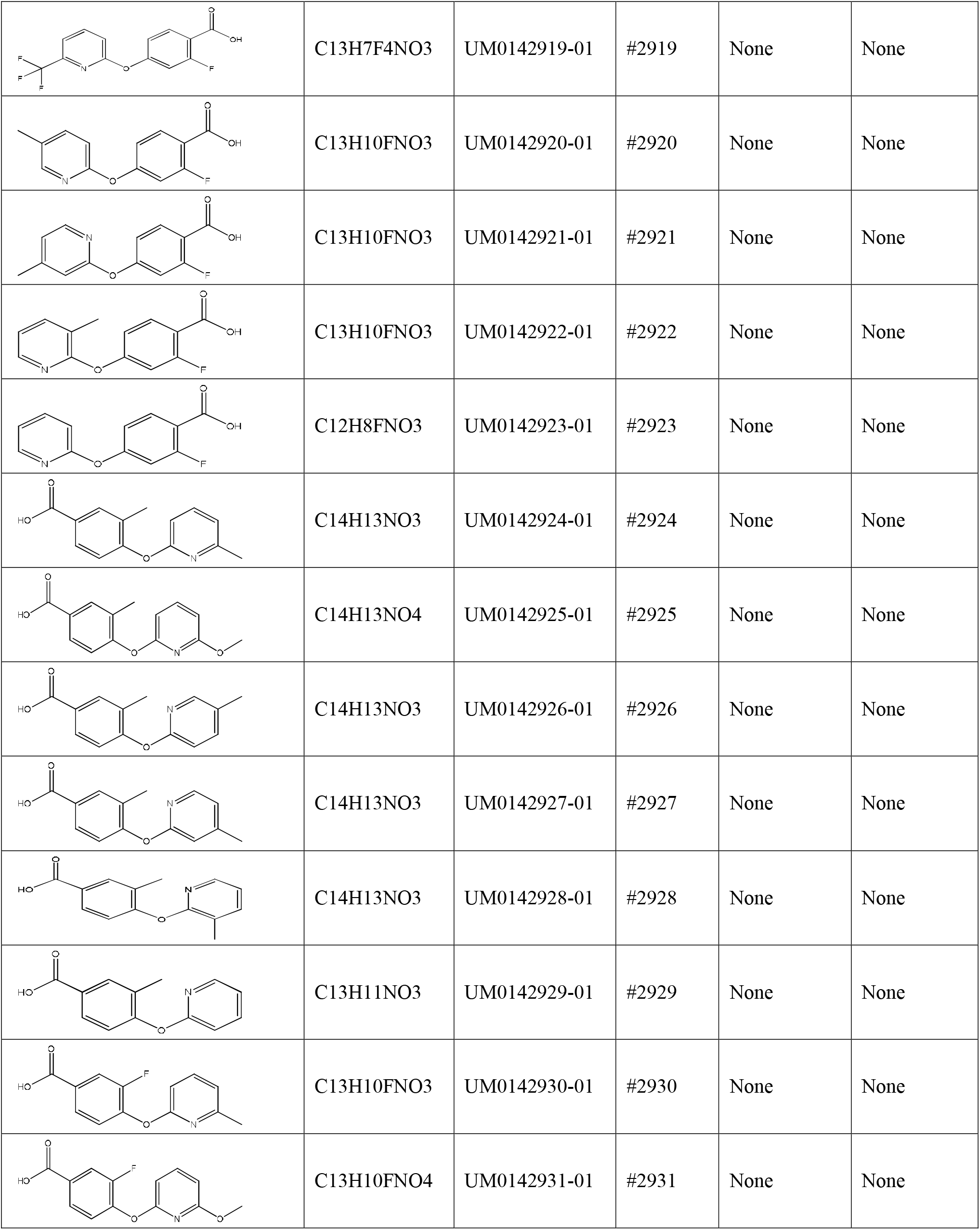
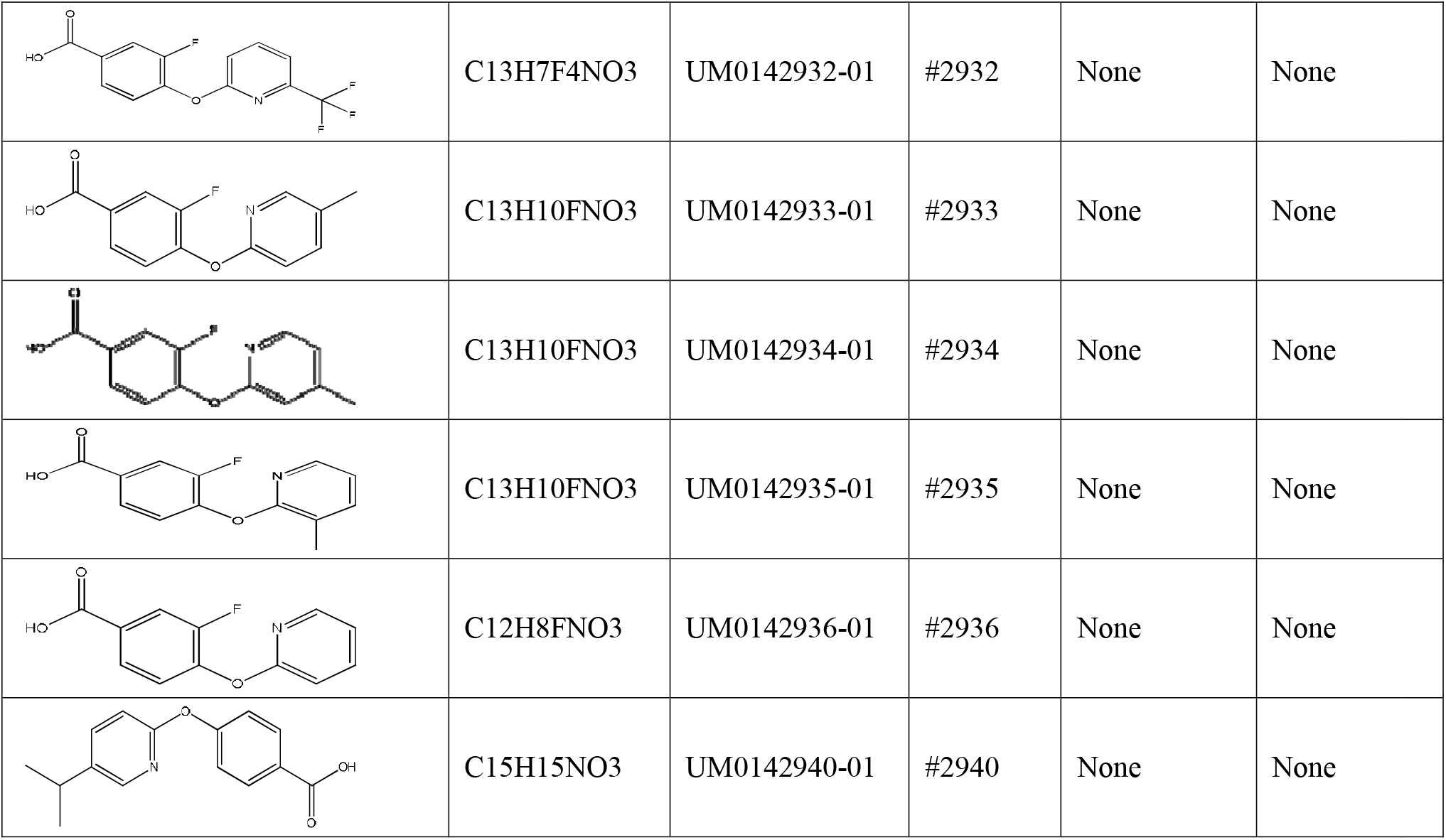
List of 1G2 derivatives created by medicinal chemistry. Structures, composition, names assigned after synthesis, numbers, AGS cell cytotoxicity and inhibition of *H. pylori* 26696 growth.

